# ESCRT-III acts in scissioning new peroxisomes from the ER

**DOI:** 10.1101/147603

**Authors:** Fred D. Mast, Thurston Herricks, Kathleen M. Strehler, Leslie R. Miller, Ramsey A. Saleem, Richard A. Rachubinski, John D. Aitchison

**Author notes:** Current address: Amgen, 360 Binney Street, Cambridge, Massachusetts, 02142-1011 (Ramsey A. Saleem). Corresponding author: John D. Aitchison, PhD, Center for Infectious Disease Research, 307 Westlake Avenue North, Suite 500, Seattle, Washington, 98109-5219.

## Abstract

Dynamic control of peroxisome proliferation is integral to the peroxisome’s many functions. A breakdown in the ability of cells to form peroxisomes is linked to many human health issues, including defense against infectious agents, cancer, aging, heart disease, obesity and diabetes, and forms the basis of a spectrum of peroxisomal genetic disorders that cause severe neuropathologies. The ER serves as a source for preperoxisomal vesicles (PPVs) that mature into peroxisomes during de novo peroxisome biogenesis and to support growth and division of existing peroxisomes. However, the mechanism of PPV formation and release from the ER remains poorly understood. Here we show that the evolutionarily ancient endosomal sorting complexes required for transport (ESCRT)-III are peroxisome biogenesis factors that function to cleave PPVs budding from the ER into the cytosol. Using comprehensive morphological and genetic assays of peroxisome formation and function we find that absence of ESCRT-III proteins impedes de novo peroxisome formation and results in an aberrant peroxisome population in vivo. Using a cell-free PPV budding assay we show that ESCRT-III proteins Vps20 and Snf7 are required to release PPVs from the ER. ESCRT-III is therefore a positive effector of membrane scission for vesicles budding both away from and towards the cytosol, a finding that has important implications for the evolutionary timing of emergence of peroxisomes and the rest of the internal membrane architecture of the eukaryotic cell.

## Introduction

Peroxisomes are membrane-bound, spherical organelles that are found across eukaryotes (Smith and Aitchison, 2013). All peroxisomes share common mechanisms guiding their biogenesis, division and protein import. The conserved protein products of *PEX* genes, termed peroxins, mediate the formation and maintenance of peroxisomes (Distel et al., 1996). However, peroxisomes in different organisms host different metabolic pathways and perform diverse functions (Mast et al., 2010). In multicellular organisms, peroxisomes perform both broad and distinct cell-type specific functions; for example, in plants peroxisomes are the sole site for β-oxidation of fatty acids and participate broadly in pathogen defense, however, in stomata they assist in stromal opening, and in mesophyll cells specialized ‘leaf’ peroxisomes are found tightly juxtaposed to chloroplasts and participate in photorespiration (Kao et al., 2018; Reuman and Weber, 2006). These examples are not exhaustive and peroxisomes perform a variety of other essential and diverse biosynthetic and catabolic processes (Reumann and Bartel, 2016).

Peroxisomes respond dramatically to different stimuli. They are induced in metazoans in response to fats, hypolipidemic agents and non-genotoxic carcinogens, and during development and differentiation (Weller et al., 2003). Peroxisome proliferation is linked to their varied functions including, for example, in response to cirrhosis of the liver (De Craemer et al., 1993), after ischemia in the brain (Young et al., 2015), in developing cardiomyocytes (Colasante et al., 2015), in protecting the auditory canal against sound-induced hearing loss from reactive oxygen species (Delmaghani et al., 2015), in epithelial cells to modulate the innate immune system (Dixit et al., 2010; Odendall et al., 2014), in macrophages to assist in clearance of microbial pathogens (Di Cara et al., 2017), and as regulatory sites for the mTOR pathway (Tripathi and Walker, 2016; Zhang et al., 2013; Zhang et al., 2015). In many of these examples, the proliferation of peroxisomes appears to be in response to metabolic need or as part of a stress response to deal with increased levels of harmful molecules, particularly reactive oxygen species, produced through insult or injury. Why this response leads to increased numbers of peroxisomes as opposed to an increase in size is unclear. For example, mice lacking the peroxisome division protein PEX11β have reduced numbers of larger, functional peroxisomes, which does not impede metabolic capacity per se, but yet these mice still display many hallmark phenotypes of peroxisome biogenesis disorder (PBD) patients and die shortly after birth (Li et al., 2002). In contrast, the link between peroxisome proliferation and the role of peroxisomes as spatial beacons for integrating and initiating signaling cascades has become clearer (Mast et al., 2015). Here, changes in peroxisome numbers contribute to tipping the balance between one signaling response versus another, such as in peroxisome-mediated innate immune signaling (Odendall et al., 2014) and regulation of the mTOR pathway (Tripathi and Walker, 2016).

Peroxisome proliferation occurs via two partially redundant mechanisms: the division of existing peroxisomes through fission, and de novo formation from the ER (Mast et al., 2015; Smith and Aitchison, 2013). Fission of peroxisomes is comparatively well characterized and requires the Pex11 family of proteins to elongate and constrict the organelle, permitting GTP-dependent scission by dynamin-related proteins (DRPs) (Schrader et al., 2016). Peroxisomes also share division components with other organelles, particularly the mitochondrion (Motley et al., 2008). In contrast, the mechanism of de novo formation remains poorly understood, and the identities of many factors involved in this process remain unknown (Agrawal and Subramani, 2016).

In yeast, biogenesis via fission dominates peroxisome proliferation (Motley and Hettema, 2007). However, most peroxisomal membrane proteins (PMPs) transit through the ER on their way to peroxisomes (Hoepfner et al., 2005; Schuldiner et al., 2008; Thoms et al., 2012; van der Zand et al., 2010). A global analysis of localized protein synthesis also revealed that many PMPs are likely cotranslated at the ER (Jan et al., 2014). Similar observations of PMP cotranslation (Kaewsapsak et al., 2017) and transport from ER to peroxisomes made in substantially divergent species of yeast (Agrawal et al., 2011; Farre et al., 2017; Titorenko and Rachubinski, 1998), plants (Hu et al., 2012), the excavate pathogen *Trypanosoma brucei* (Bauer et al., 2017; Guther et al., 2014), and mammalian cells (Kim et al., 2006; Mayerhofer et al., 2016) highlight the essential and evolutionarily ancient role of the ER in peroxisome biogenesis in eukaryotic cells.

A vesicular transport pathway transfers proteins and membranes from the ER to peroxisomes (Agrawal et al., 2016; Agrawal et al., 2011; Lam et al., 2010). This pathway is essential even when peroxisomes multiply by growth and division, as altering the flux through this transport pathway alters the numbers of peroxisomes in cells (Mast et al., 2016). Pex3 accumulates initially at an ER subdomain before being released in a preperoxisomal vesicle (PPV) that buds from the ER (Halbach et al., 2006; Hoepfner et al., 2005; Tam et al., 2005). Sorting of PMPs through the ER to sites of PPV formation and egress requires both Pex3-dependent and-independent processes (Fakieh et al., 2013). The ER-shaping reticulon proteins, through physical interaction with Pex29 and Pex30, assist in regulating Pex3 sorting through the ER and releasing PPVs (David et al., 2013; Mast et al., 2016). Pex30 and its paralogue, Pex31, have membrane-shaping capabilities like the reticulon proteins, which may help in defining and segregating the PPV exit site in the ER (Joshi et al., 2016).

The formation of PPVs requires Pex3 and Pex19, and loss of PPV formation leads to a cell’s inability to form peroxisomes and consequently its eventual loss of the organelle (Hettema et al., 2000; Hoepfner et al., 2005). Pex19 is a cytosolic protein that interacts with Pex3 and other PMPs (Agrawal et al., 2017), functioning as a chaperone, and is essential for budding PPVs from the ER (Lam et al., 2010). At least two classes of PPVs (V1 and V2) have been characterized (Agrawal et al., 2016; Titorenko et al., 2000; Titorenko and Rachubinski, 2000; van der Zand et al., 2012); both contain Pex3 but differ in whether they contain or lack docking-factor or RING finger group proteins of the peroxisomal matrix protein import complex (peroxisomal importomer) (Agrawal et al., 2016). The separation of these two subcomplexes could prevent premature assembly of the peroxisomal importomer in the ER and the potential import of peroxisomal matrix proteins directly into the ER (Agrawal et al., 2016; van der Zand et al., 2012). While Pex3 and Pex19 are necessary for PPV budding, they are not sufficient for this process, and evidence suggests additional cytosolic component(s) are required, at least one of which likely consumes ATP (Agrawal et al., 2011; Lam et al., 2010). DRPs, which function in peroxisome division, or COPI and COPII vesicle transport pathways, all of which consume GTP, have repeatedly been shown not to be required for PPV formation (Lam et al., 2010; Motley et al., 2015; Motley and Hettema, 2007; Perry et al., 2009; South et al., 2000).

Here, we report a novel role for endosomal sorting complexes required for transport (ESCRT)-III in the de novo biogenesis of peroxisomes. In particular, using a series of comprehensive morphological and genetic assays of peroxisome formation and function and in vitro biochemical assays that produce PPVs from the ER, we implicate ESCRT-III as being essential for the scission of PPVs from the ER.

## Results

### Genomic screens identify a putative role for ESCRT in peroxisome biogenesis

Screens of an isogenic, arrayed collection of yeast gene deletion strains identified 211 genes whose disruption led to defects in the cell’s ability to form peroxisomes (Saleem et al., 2010). We reasoned that this dataset held clues to candidates involved in the de novo biogenesis of peroxisomes and in the formation of PPVs at the ER. Candidates would be expected a priori to be cytosolic and/or localized to the ER, and to use ATP in their activity. Components of ESCRT exhibit these characteristics and were enriched in this dataset with a statistically significant hypergeometric p-value of 0.003 (Saleem et al., 2010).

ESCRT is composed of five subcomplexes, including ESCRT-0,-I,-II,-III (and-III-associated), and the AAA-ATPase Vps4 complex (Babst et al., 2002a; Babst et al., 2002b; Babst et al., 1997; Katzmann et al., 2001; Katzmann et al., 2003) (reviewed in (Henne et al., 2011; Schoneberg et al., 2016)). These five complexes function sequentially to mediate the formation of intraluminal vesicles and also assist in piecemeal fashion in numerous other cellular activities like cytokinesis (Carlton and Martin-Serrano, 2007), plasma membrane repair (Jimenez et al., 2014; Scheffer et al., 2014), autophagosome closure (Lee et al., 2007; Rusten et al., 2007), viral replication and budding (Garrus et al., 2001), nuclear envelope reformation (Olmos et al., 2015; Vietri et al., 2015), nuclear pore complex quality surveillance (Webster et al., 2014), neuronal pruning (Loncle et al., 2015), and microtubule severing (Guizetti et al., 2011) (Henne et al., 2011; Schoneberg et al., 2016). ESCRT-III, composed of Vps20, Snf7, Vps24 and Did4 in yeast, is evolutionarily ancient and is the primary effector complex for all these activities (Henne et al., 2011; Tang et al., 2015). Given this wealth of function, particularly for ESCRT-III, a key consideration was whether ESCRTs have a direct role in peroxisome biogenesis or exert indirect effects due to their diversity of activity. In our global screens for peroxisome effectors (Saleem et al., 2010; Smith et al., 2006), ESCRT deletion strains showed defects in metabolizing fatty acid carbon sources, a process requiring functional peroxisomes; had altered expression of the peroxisomal β-oxidation enzyme 3-ketoacyl-CoA thiolase (Pot1); but exhibited only mild or no peroxisome morphology defects as assessed by high-throughput fluorescence microscopy (Table S1).

### Yeast strains lacking ESCRT-III components exhibit oleic acid-specific growth defects

We used our one-cell doubling evaluation of living arrays of yeast (ODELAY!) platform (Herricks et al., 2017a) to re-address the role of ESCRT in peroxisome biogenesis by comparing growth on solid-phase medium containing as the sole carbon source either glucose or oleic acid, which requires functional peroxisomes for its metabolism (Figs. 1 and S1). Unlike traditional spot-based assays or assays using optical density measurements at a population level, ODELAY! provides time-resolved measurements of yeast populations growing from individual cells into colonies, for an array of up to 96 different strains (Herricks et al., 2017a). ODELAY! therefore permits standardized analyses of the growth rates and population heterogeneity both within and between strains measured in parallel across different growth conditions. Strains with general growth defects have comparable normalized growth rates under both conditions, whereas condition-specific strains grow more slowly under one condition versus another. We measured growth rates for all 18 ESCRT deletion strains, 6 peroxin/peroxisome-related control deletion strains, and 20 randomly selected deletion strains from the yeast deletion library (Fig. 1 A and Table S2).

**Figure 1.**
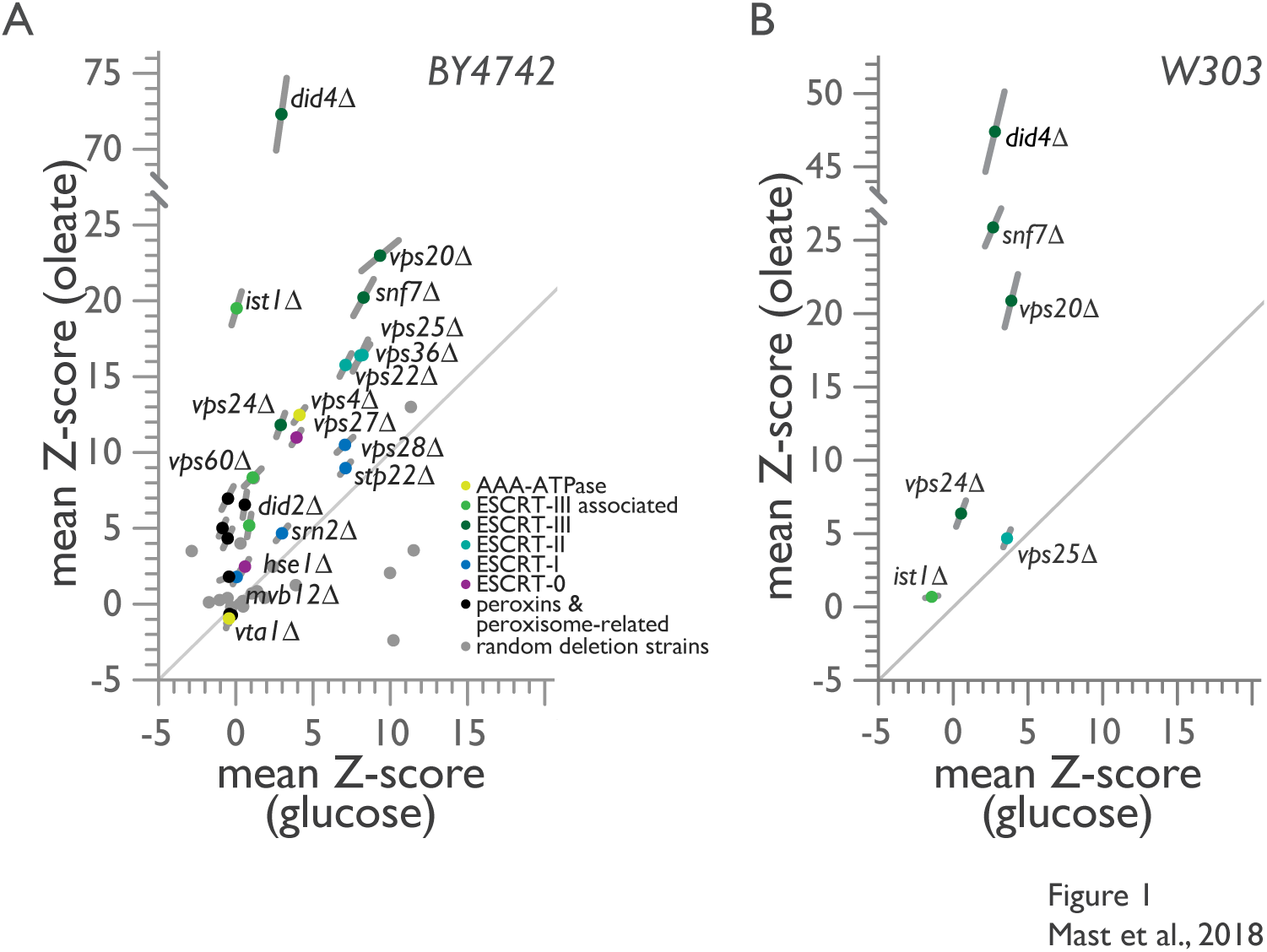
ESCRT-III mutants exhibit oleate-specific growth defects. Doubling times of individual yeast cells growing into colonies on solid-phase medium containing either glucose or oleic acid as sole carbon source were measured with ODELAY!. A population-derived *Z*-score and SEM for each strain are plotted with measurements from glucose-containing medium on the *x*-axis against measurements from oleic acid-containing medium on the *y*-axis from 8 biological replicates per growth condition. (A) ODELAY! results for deletion strains derived from the *BY4742* parental strain. (B) ODELAY! results for select ESCRT deletion strains derived from the *W303* parental strain.

Strain doubling times measured by ODELAY! were normalized, and the mean *Z*-score for each deletion strain was compared between the two growth conditions (Fig. 1 A). The *pex19*Δ, *pex13*Δ and *pex14*Δ strains, which lack the ability to form peroxisomes (Hettema et al., 2000) and a functional peroxisomal importomer (Meinecke et al., 2010), or the *pot1*Δ strain lacking the enzyme that performs the last step of peroxisomal β-oxidation (Igual et al., 1991), revealed condition-specific growth defects, as they exhibited slightly faster or marginally slower growth in the presence of glucose but grew 4-5 standard deviations more slowly than the corresponding wild-type strain *BY4742* in the presence of oleic acid (Fig. 1 A).

ESCRT-III deletion strains showed pronounced condition-specific growth defects in the presence of oleic acid, with *did4*Δ cells registering the slowest doubling rate of all ESCRT deletions, followed by *vps20*Δ, *snf7*Δ and *ist1*Δ, which is a deletion strain for an ESCRT-III associated gene (Fig. 1 A). ESCRT-I deletion strains displayed general growth defects, whereas ESCRT-II deletion strains clustered tightly, in line with their known structural and functional assembly as a single protein complex, and showed a condition-specific growth defect in oleic acid, although not as severe as that observed for strains deleted for ESCRT-III components (Fig. 1 A). Other ESCRT deletion strains also had slower, but less severe, doubling times in oleic acid than ESCRT-III deletion strains, except for the strain lacking the Vps4 regulator Vta1 which, like the peroxisome inheritance mutants *inp1*Δ and *inp2*Δ, had a faster doubling time than the wild-type strain under both glucose and oleic acid conditions (Fig. 1 A).

Despite the skew to oleic acid-specific growth defects observed for the majority of ESCRT deletion strains in the *BY4742* background, the randomly selected gene deletions revealed no bias in our assay (Fig. 1 A and Table S2). Of note, *sec28*Δ, a strain lacking the ɛ-COP subunit of coatomer (Duden et al., 1998), demonstrated a significant growth defect when grown on medium containing glucose but grew significantly faster than the wild-type strain on medium containing oleic acid (Fig. 1 A and Table S2). This phenotype could partially result from the overall slower growth of cells metabolizing oleic acid, which would mask defects in COPI vesicle trafficking, and also further demonstrates that COPI is not involved in peroxisome biogenesis. Other deletion strains affecting the secretory pathway, such as *emp46*Δ, lacking a COPII-associated protein (Sato and Nakano, 2002), or *nyv1*Δ, lacking a v-SNARE of the vacuolar SNARE complex (Nichols et al., 1997), showed negligible general growth defects (Fig. 1 A and Table S2). Therefore, general defects in endocytic trafficking do not manifest as condition-specific defects in the metabolism of the non-fermentable carbon source, oleic acid.

We repeated our ODELAY! analysis in the *W303* strain background and evaluated strains deleted for the genes *VPS20, SNF7, DID4* and *VPS24* encoding ESCRT-III components, together with strains deleted for the genes *IST1* and *VPS25* (Fig. 1 B and Table S2). We observed similar condition-specific growth defects for the ESCRT-III deletion strains in the presence of oleic acid in the *W303* strain background as we had observed in the *BY4742* strain background (Fig. 1 A and Table S2) However, for *vps25*Δ, while oleic acid-specific defects were observed in the *BY4742* strain deleted for *VPS25* (Fig. 1 A), these condition-specific defects were not reproduced in the *W303* strain deleted for *VPS25*, which displayed general and equivalent growth defects on both glucose and oleic acid carbon sources (Fig. 1 B). The *ist1*Δ strain, which showed condition-specific growth defects in *BY4742*, showed negligible growth defects in *W303*, where it grew faster than *wild-type* in the presence of glucose and slightly slower than *wild-type* in the presence of oleic acid (Fig. 1 B). We therefore conclude that ESCRT-III has a specific functional role in the metabolism of the non-fermentable carbon source, oleic acid.

### Loss of ESCRT-III components results in peroxisomes with aberrant morphologies

To determine if the growth defects observed for ESCRT-III mutants result from defects in peroxisomes, we used EM to investigate the cellular ultrastructure of cells lacking individual components of the core ESCRT-III complex under conditions that promote peroxisome biogenesis (Figs. 2 and S1). Yeast were grown in the presence of oleic acid for 8 h to induce peroxisome proliferation and peroxisomal membrane expansion (Saleem et al., 2008; Smith et al., 2002). Under these conditions peroxisomes were readily observed in *wild-type* cells as orbicular structures delimited by a single lipid bilayer and containing an electron-dense, paracrystalline matrix (Fig. 2 A). In contrast, peroxisomes were not observed in *vps20*Δ or *snf7*Δ cells, which instead contained infrequent small vesicular structures of unknown origin, although often in close apposition to, and possibly contiguous with, ER membranes (Fig. 2, B and C). Quantification revealed fewer peroxisomes and vesicular-like structures for all deletion strains of the ESCRT-III complex (Fig. 2 D and Table 1), although peroxisomes similar in size to *wild-type* peroxisomes were observed in *did4*Δ and *vps24*Δ cells, but at levels 25% and 45%, respectively, of peroxisomes in *wild-type* cells (Fig. 2, E - I).

**Figure 2.**
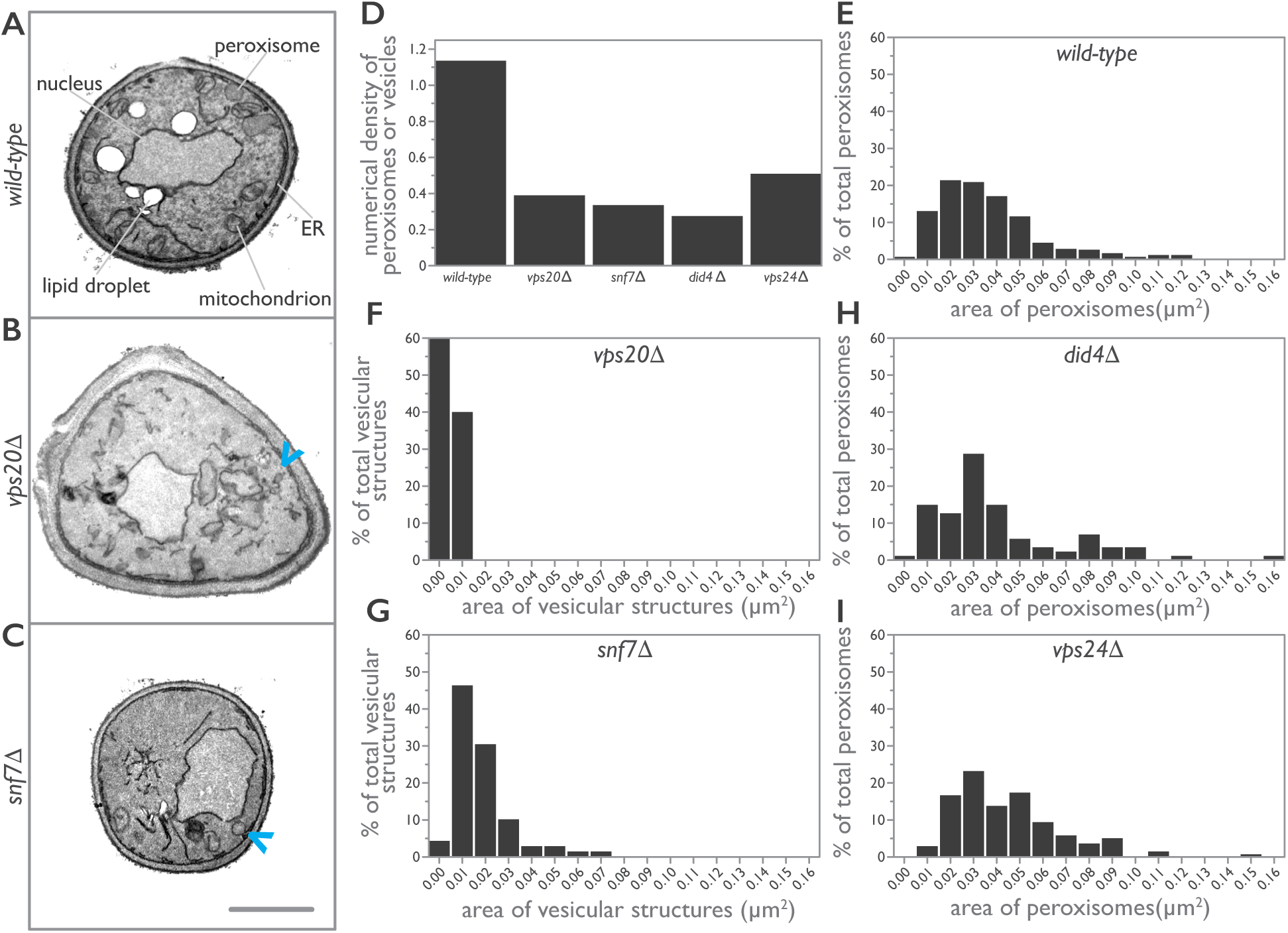
ESCRT-III deletion strains have populations of smaller and aberrant peroxisomes. (A) *Wild-type*, (B) *vps20*Δ and (C) *snf7*Δ cells were grown in medium containing oleic acid for 8 h to induce the production of peroxisomes and prepared for EM. Representative cell ultrastructures and organelle profiles are presented. Arrowheads in (B) and (C) point to aberrant vesicular structures observed in *vps20*Δ and *snf7*Δ cells. Bar, 1 μm. (D–I) Morphometric analysis of peroxisomes. For each strain analyzed, the areas of individual peroxisomes were determined and used to calculate the numerical density of peroxisomes, which is the number of peroxisomes per μm^3^ of cell volume (Weibel and Bolender, 1973). Bar graphs display the numerical density of peroxisomes (D) and the population distribution of the areas of peroxisomes (E–I) for *wild-type* and ESCRT-III deletion strains.

**Table 1.** Average area and numerical density of peroxisomes or vesicular structures in cells of *wild-type* and ESCRT-III deletion strains

| strain | cell area<br>assayed ($\mu\text{m}^2$) | peroxisome<br>count | numerical density of<br>peroxisomes<sup>#</sup> | average area of<br>peroxisomes ($\mu\text{m}^2$) |
| --- | --- | --- | --- | --- |
| *BY4742* | 1396.09 | 421 | 1.14 | 0.037 |
| *vps20Δ* | 1125.46 | 69* | 0.39* | 0.005* |
| *snf7Δ* | 1144.83 | 89* | 0.34* | 0.018* |
| *did4Δ* | 820.13 | 35 | 0.28 | 0.040 |
| *vps24Δ* | 933.70 | 138 | 0.51 | 0.044 |

\* denotes vesicular structures, as no peroxisomes were observed in these strains.

### ESCRT-III components act as positive effectors of de novo peroxisome assembly

The observed growth and peroxisome morphology defects detected for deletion strains of ESCRT-III and in particular for *vps20*Δ and *snf7*Δ mutants suggested a role for ESCRT-III in peroxisome biogenesis. To test this, we used an in vivo peroxisome biogenesis assay in which tetracycline control of *PEX19* expression permits synchronization of cells for de novo peroxisome biogenesis without overexpressing peroxins (Fig. 3 A) (Mast et al., 2016). This is a critical distinction from de novo biogenesis assays that rely on repression and then overexpression of a peroxin, as it does not lead to mistargeting of the peroxisomal reporter to the general ER or other organelles. With *PEX19* expressed, *wild-type* cells contained an average of 10 peroxisomes per cell, as assessed by the punctate localization of peroxisomal Gpd1-GFP (Fig. 3 A) (Jung et al., 2010). Repression of *PEX19* expression by treatment with doxycycline (DOX) for 24 h removed peroxisomes from the *wild-type* yeast population, with only ^~^0.2% of cells containing on average one peroxisome per cell. Removal of DOX enabled the expression of *PEX19* and initiated de novo biogenesis of peroxisomes (Fig. 3 A). We tracked peroxisome formation using three-dimensional fluorescence microscopy over 24 h, sampling cells at 0, 4, 8, 12 and 24 h after initiation of de novo peroxisome biogenesis. Quantification of the number of peroxisomes per cell revealed a population distribution of steady-state peroxisome densities comparable to those we have measured previously (Fig. 3 B) (Mast et al., 2016).

**Figure 3.**
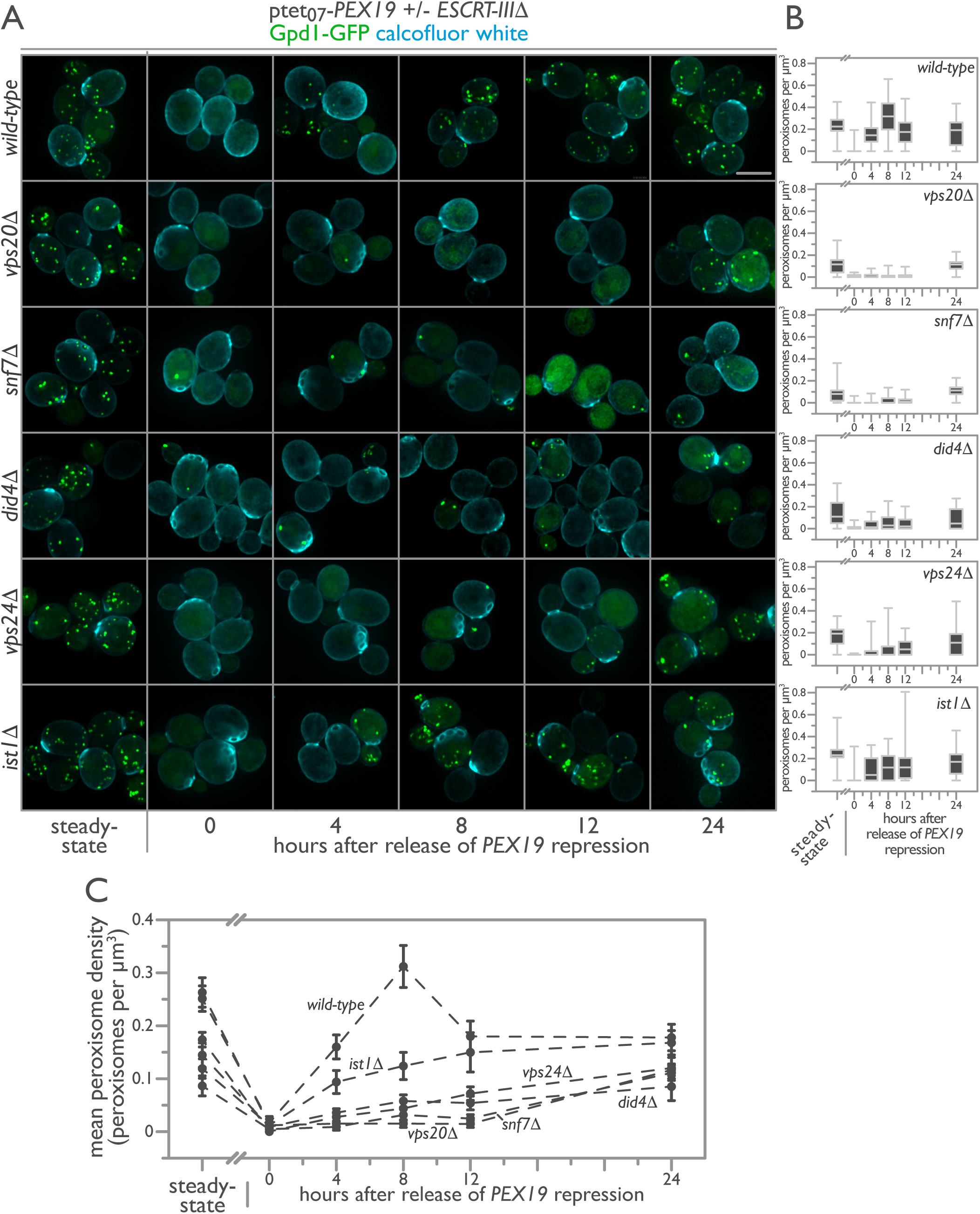
ESCRT-III positively regulates de novo peroxisome biogenesis. (A) Genomically encoded *PEX19* was placed under the control of a tetracycline-repressible tet_07_ promoter to allow regulatable de novo peroxisome production in *wild-type, vps20*Δ, *snf7*Δ, *did4*Δ, *vps24*Δ, and *ist1*Δ cells expressing Gpd1-GFP as a peroxisomal marker. Cells were imaged before (steady-state) and after 24 h incubation with 4 μM DOX. Additional observations were made 4, 8, 12, and 24 h after removal of DOX. Scale bar, 5 μm. (B) The population distributions of peroxisome densities per cell are depicted at each time point as interquartile box-and-whisker blots. (C) The mean peroxisome density per strain over a time course of de novo peroxisome biogenesis was plotted with error bars representing the SEM of 3 biological replicates.

The assembly of peroxisomes de novo is a dynamical systems process that proceeds non-linearly following reintroduction of *PEX19* (Fig. 3). In *wild-type* cells, peroxisomes reappeared in 91% of cells 4 h after induction of *PEX19* and proceeded to overshoot steady-state levels of peroxisomes 8 h post-induction. By 12 h, the population distribution of peroxisome levels had dropped below those at steady-state, and by 24 h were still on a trajectory to return to steady-state levels. The dynamics of de novo peroxisome formation were strongly influenced by the length of repression of *PEX19* and by the state of cell growth, in accordance with our previous observations (Mast et al., 2016). These current observations imply the existence of positive and negative regulators of de novo peroxisome biogenesis.

Strikingly, ESCRT-III mutants displayed severe defects in de novo peroxisome biogenesis. ESCRT-III mutants had fewer peroxisomes per cell under steady-state conditions and retained more peroxisomes per cell following DOX repression of *PEX19* (Fig. 3). Upon stimulation of de novo peroxisome biogenesis, *vps20*Δ, *snf7*Δ, *did4*Δ and *vps24*Δ cells, in order of severity, failed to produce peroxisomes at rates comparable to that of *wild-type* cells, revealing a defect in the ability of *vps20*Δ and *snf7*Δ cells especially to form new peroxisomes de novo (Fig. 3).

Pex19 dynamics in *wild-type* cells presaged peroxisome dynamics in these cells, as Pex19 levels returned to steady-state 4 h after release of DOX-induced repression, then leveled off and dropped slightly at 8 and 12 h, before increasing slightly at 24 h post-induction (Fig. 4). Like *wild-type*, ESCRT-III mutants showed no detectable levels of Pex19 following DOX treatment, but by 4h post-induction Pex19 levels were comparable to those found in *wild-type* cells. However, Pex19 dynamics were altered in the ESCRT-III mutants as compared to *wild-type* (Fig. 4). We conclude that while Pex19 dynamics were altered in ESCRT-III mutants, ESCRT-III mutants nonetheless expressed sufficient Pex19 at all times following removal of DOX to rule out a contribution of altered Pex19 dynamics as the primary explanation for the catastrophic de novo biogenesis defects observed in these strains.

**Figure 4.**
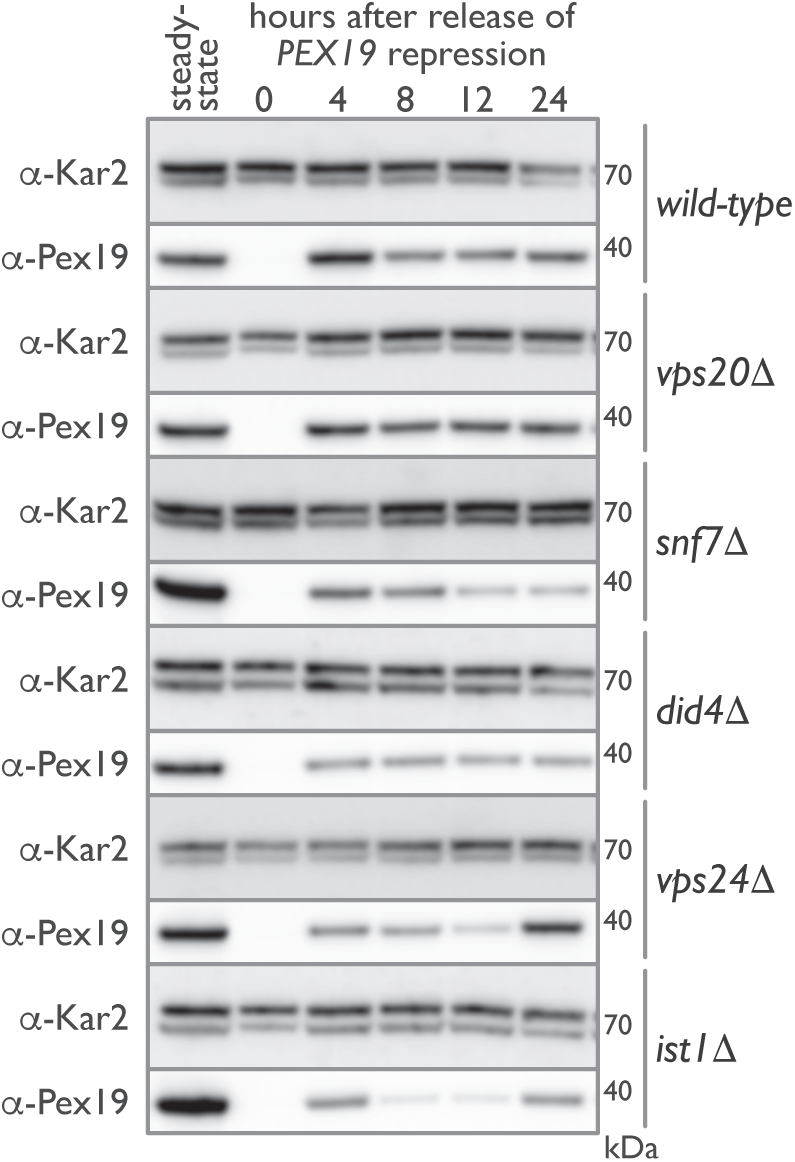
Pex19 dynamics are altered in ESCRT-III deletion mutants during de novo peroxisome biogenesis. Genomically encoded *PEX19* was placed under the control of a tetracycline-repressible tet_07_ promoter to allow regulatable de novo peroxisome production in *wild-type, vps20*Δ, *snf7*Δ, *did4*Δ, *vps24*Δ and *ist1*Δ cells. Whole cell lysates were prepared from each strain before (steady-state) and after a 24 h incubation with 4 μM DOX. Additional whole cell lysates were prepared 4, 8, 12 and 24 h after removal of DOX. Equal amounts of protein from these whole cell lysates were resolved by SDS-PAGE, transferred to PVDF, and immunoblotted for Pex19 and Kar2, which was used as a control for protein loading.

### ESCRT-III functions conditionally downstream of Pex19

The interplay between Pex19 dynamics and peroxisome dynamics and the interruption of that interplay in ESCRT-III mutants suggested to us the potential for genetic interactions between Pex19 and ESCRT-III. Genetic interactions can often provide insights into the function of proteins, order pathways, and reveal coordinated functions (Costanzo et al., 2016; Schuldiner et al., 2005). Common examples of genetic interactions include epistasis, where two mutants have phenotypes different from that of *wild-type* and the double mutant shows the phenotype of one of the single mutants. These “alleviating” interactions suggest that one protein functions upstream of the other in a linear pathway (Mani et al., 2008). Similarly, synthesis occurs when the phenotype of a double mutant is more severe than an additive effect of the individual mutant phenotypes. Common interpretations of this scenario include proteins that function in parallel biological pathways or form structures that cannot withstand, or ‘buffer’, the loss of both components (Dixon et al., 2009). More complex modes of genetic interaction also exist, including single-nonmonotonic, where a single mutant shows opposing phenotypes in the *wild-type* and other single mutant backgrounds (Drees et al., 2005). There are no common interpretations for this scenario, but such an interaction would imply that epistasis exists under a limiting set of circumstances.

Using ODELAY!, we measured growth rates for the tetracycline-regulated *PEX19* background strain as well as strains additionally harboring ESCRT-III deletion mutants under conditions where *PEX19* was differentially expressed. The normalized mean *Z*-scores were plotted on a phenotypic axis for each ESCRT-III mutant alongside its corresponding phenotype (Φ) inequalities (Fig. 5) (Drees et al., 2005). This unbiased and quantitative analysis revealed complex genetic interactions between *PEX19* and ESCRT-III with *did4*Δ, *vps20*Δ and, to a lesser extent, *snf7*Δ being single-nonmonotonic to *PEX19* depletion. Whereas *vps24*Δ, and potentially *snf7*Δ, is epistatic to *PEX19* depletion, *ist1*Δ is double-nonmonotonic with *PEX19* depletion. We interpret these results to suggest that ESCRT-III functions conditionally downstream of *PEX19*, i.e. the function of ESCRT-III is contextual and specific to the budding of peroxisomes from the ER.

**Figure 5.**
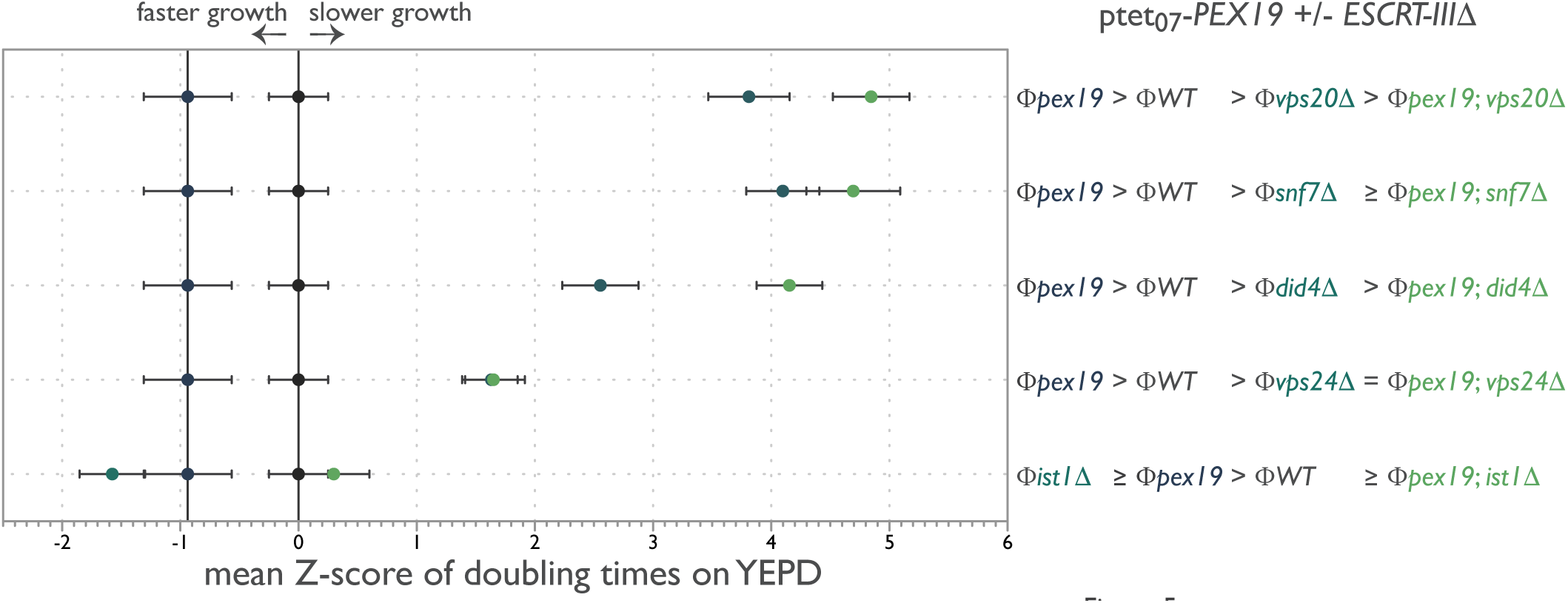
ESCRT-III functions conditionally with *PEX19*. Doubling times of individual yeast cells growing into colonies on solid-phase YEPD medium with and without 4 μM DOX were measured with ODELAY!. Strains grown in the presence of DOX were precultured for 24 h in the presence of 4 μM DOX to turn off *PEX19* and deplete Pex19 levels from cells. A population-derived *Z*-score and SEM for each strain are plotted on a phenotypic axis and summarized as a phenotypic inequality defining the genetic interaction between the indicated genes from 4 biological replicates per strain per growth condition.

Notably, ESCRT-III localized to sites of de novo peroxisome biogenesis. As expected, Pex3-GFP and Snf7-mCherry puncta rarely overlapped under steady-state control conditions, but 13.5% of Pex3p-GFP puncta correspondingly colocalized with the Snf7-mCherry signal following release from DOX treatment to induce de novo peroxisome formation (Fig. 6 A). Interestingly, Pex3-GFP foci colocalizing with Snf7-mCherry were dimmer and sometimes elongated, similar in appearance to previous observations of Pex3 in cells repressed for members of the *DSL1* protein complex (Perry et al., 2009) and for Pex3 that colocalized with Pex30 and Pex29 in the ER under steady-state conditions (Mast et al., 2016). We were unable to detect a meaningful fluorescence signal for Vps20-mCherry and Vps24-mCherry under these conditions; however, Did4-mCherry showed a pattern similar to that of Snf7-mCherry despite having a very low signal-to-noise ratio (Fig. 6 B). The reduction in colocalization of Did4-mCherry compared to Snf7-mCherry may be biologically relevant or may reflect the low fluorescence signal for Did4-mCherry that was measured for these cells.

**Figure 6.**
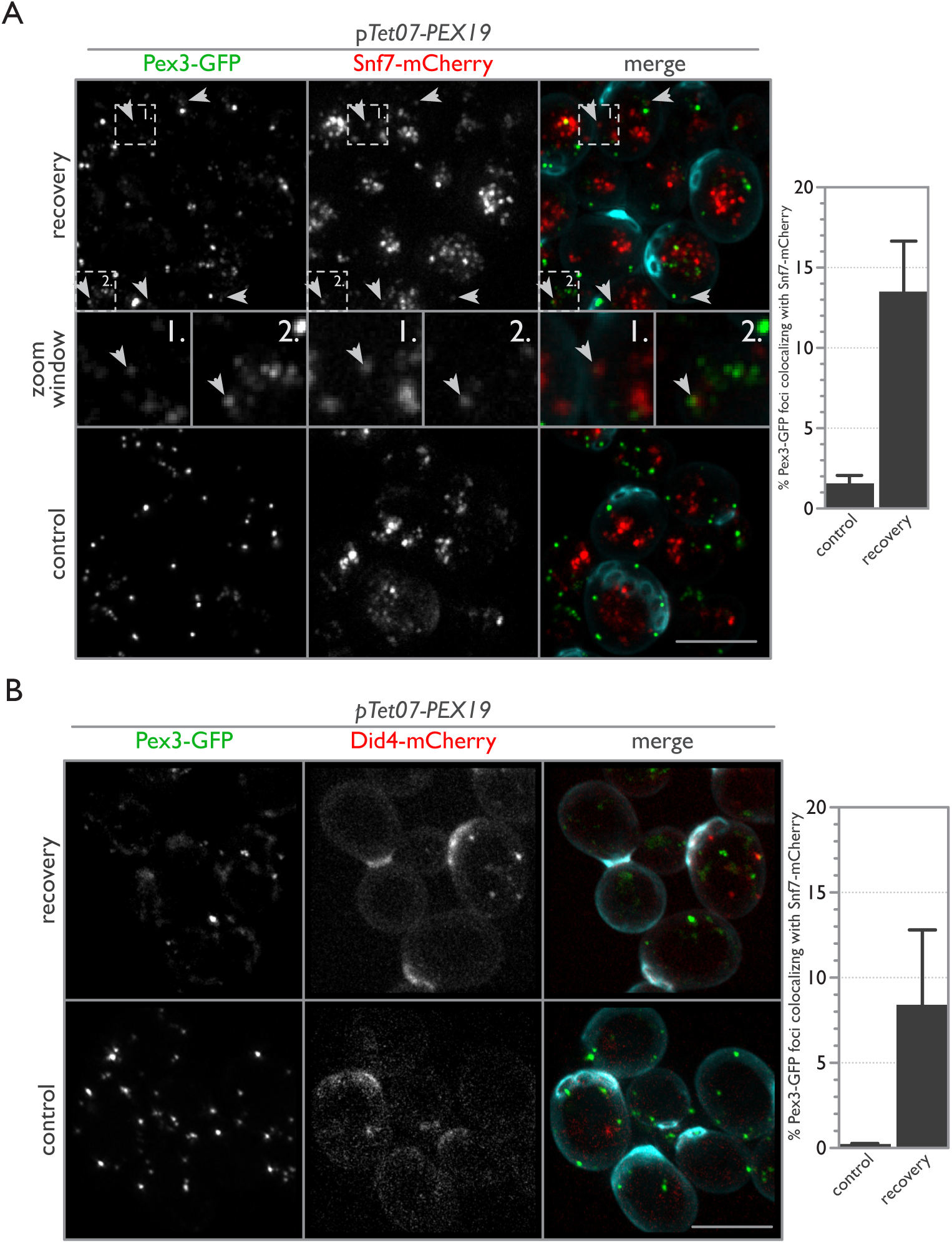
ESCRT-III components localize to sites of de novo peroxisome biogenesis. (A) Snf7 localizes to sites of de novo peroxisome biogenesis. *wild-type* cells expressing endogenously tagged Pex3-GFP and Snf7-mCherry with tetracycline-repressible *PEX19* were imaged before (control) and 1 h following an 18 h incubation with 2 μM DOX (recovery). Arrowheads point to areas of colocalization between Pex3-GFP and Snf7-mCherry (see zoom window for enlargement). For the merged image, calcofluor white staining (cyan) was used to demarcate cell boundaries. Bar graphs report the percent overlap of Pex3-GFP puncta colocalizing with Snf7-mCherry puncta with error bars representing the SEM of 3 biological replicates. Scale bar, 5 μm. (B) Same as (A), but with Did4-mCherry instead of Snf7-mCherry.

### ESCRT-III is required for the release of preperoxisomal vesicles from the ER

Current models of ESCRT-III function propose that Vps20 is recruited to a membrane first, initiating activation and polymerization of additional members, especially Snf7, to form spirals and coils that deform membranes into tubes or cones to achieve membrane scission (Henne et al., 2011; Schoneberg et al., 2016). Vps24 caps the polymers and stops polymerization, whereas Did4 is integrated into the polymer to promote its disassembly by recruiting the AAA-ATPase, Vps4 (Henne et al., 2011; Schoneberg et al., 2016).

We used an in vitro budding assay that reconstitutes the packaging and release of PMPs from the ER via PPVs (Lam et al., 2010, Mast et al., 2016) to test if ESCRT-III is required for PPV scission at the ER (Fig. 7). ER membranes were prepared from a *pex19*Δ strain in which Pex3 and other PMPs are trapped in the ER (Agrawal et al., 2016; Agrawal et al., 2011). The release of Pex3 was stimulated by addition of *wild-type* cytosol and an ATP regeneration system (Lam et al., 2010) but not by cytosol from *pex19*Δ, *vps20*Δ, or *snf7*Δ cells (Fig. 7 A and C). We also tested cytosols from other ESCRT and ESCRT-III deletion strains, e.g. *did4*Δ and *vps24*Δ, and found that their cytosols still promoted the formation of Pex3-containing PPVs but sometimes at levels significantly less than that produced by *wild-type* cytosol (Figs. 7 A and S2). Defects in budding were independent of the levels of Pex19 in ESCRT-III mutant strains (Fig. S3). The defect in PPV budding was also independent of the background yeast strain, as similar results were obtained using cytosols obtained from deletion strains in the *W303* strain background (Fig. 7 B). Mixing the 100 k*g* supernatant fraction of cytosols (S100) from *snf7*Δ, *vps20*Δ and *pex19*Δ strains pairwise in a 1:1 ratio complemented the defect in Pex3 release (Fig. 7, C and D).

**Figure 7.**
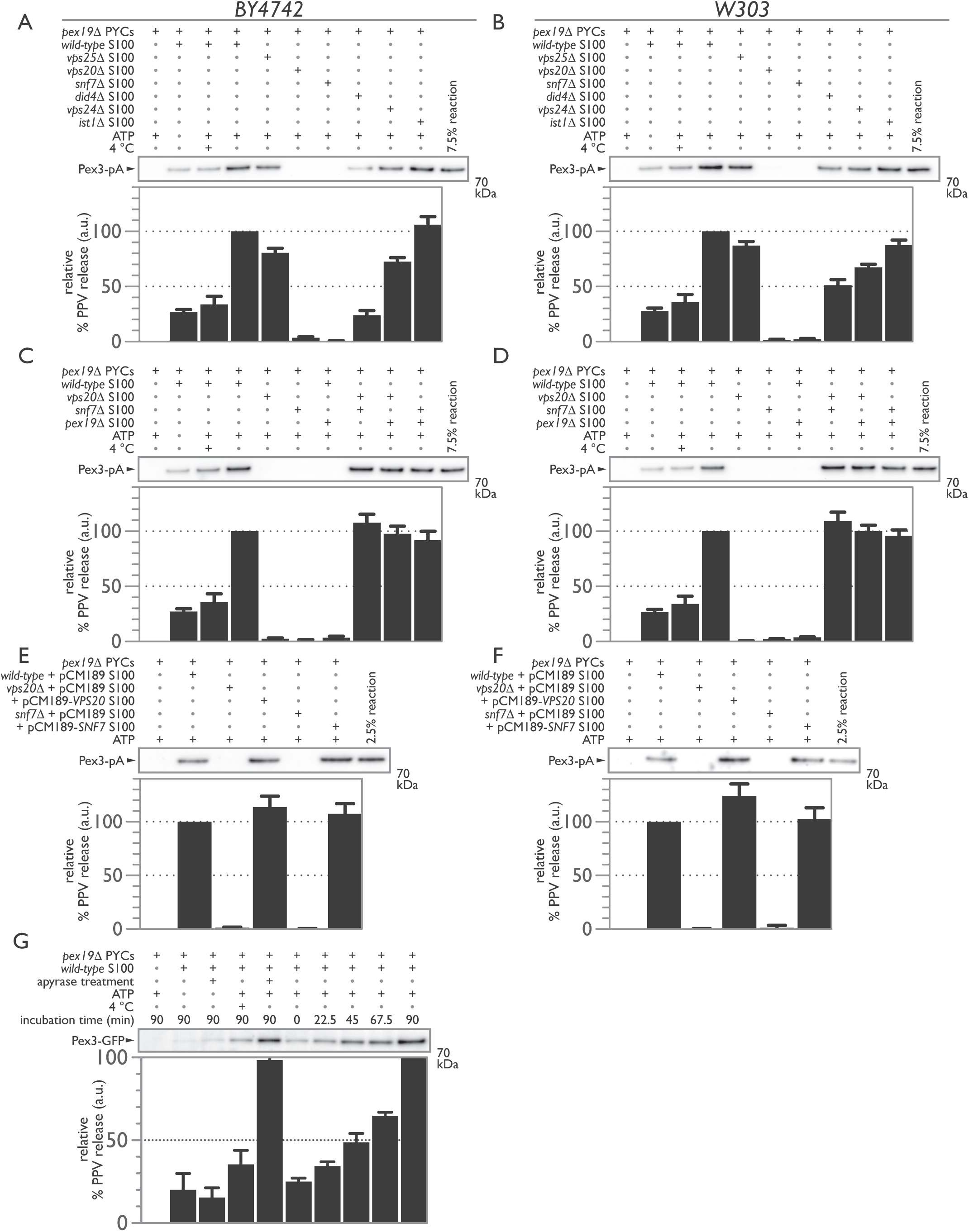
ESCRT-III is required for PPV budding from the ER. (A) Permeabilized *pex19*Δ yeast cells (PYCs) containing microsomes and expressing Pex3-pA were incubated with the S100 fraction of cytosol isolated from *wild-type* (lanes 2-4), *vps25*Δ (lane 5), *vps20*Δ (lane 6), *snf7*Δ (lane 7), *did4*Δ (lane 8), *vps24*Δ (lane 9) or *ist1*Δ (lane 10) for 90 min at room temperature in the presence of an ATP-regenerating system. Controls included incubating the PYCs alone (lane 1), with cytosol but no ATP (lane 2), or with cytosol and ATP but at 4°C (lane 3). (B) As in (A), but with the S100 fraction of cytosol isolated from *W303 wild-type* and deletion strains. (C and D) Deficiencies in PPV budding are complemented by mixing the S100 fractions of defective cytosols. PYCs were incubated with *wild-type* (lanes 2-4), *vps20*Δ (lanes 5, 8, and 9), *snf7*Δ (lanes 6, 8, and 10), and *pex19*Δ (lanes 7, 9, and 10) S100 fractions of cytosol for 90 min at room temperature in the presence of an ATP-regenerating system. Controls included incubating the PYCs alone (lane 1), with cytosol but no ATP (lane 2), or with cytosol and ATP but at 4°C (lane 3). (E and F) *VPS20* and *SNF7* complement deficiencies in PPV budding in their respective deletion mutants. PYCs were incubated with S100 fractions of cytosol from *wild-type* carrying pCM189 (lane 2), *vps20*Δ carrying pCM189 (lane 3) or pCM189-VPS20 (lane 4), or *snf7*Δ carrying pCM189 (lane 5) or pCM189-*SNF7* (lane 6). (G) ATP is not required for PPV release but rather to recycle scission components. PYCs expressing Pex3-GFP and the S100 fraction of cytosol isolated from *wild-type* were pretreated with apyrase before starting the reaction (lanes 3 and 5). The reaction was also sampled at the indicated times (lanes 6-10). For each panel, bar graphs represent the relative % of PPV release in arbitrary units with error bars representing the SEM of 3 biological replicates.

Complementation was also achieved by expressing exogenous copies of *VPS20* or *SNF7* in their respective deletion strains (Fig. 7, E and F). Here, the likely overexpression of *VPS20* led to enhanced PPV release, whereas the likely overexpression of *SNF7* did not. The production of PPVs at 4 °C suggested that scission is mechanical and not reliant on ATP-dependent enzymatic activity (Fig. 7 A – D). It is known that ESCRT-III does not depend on ATP hydrolysis to mediate membrane scission but rather relies on the AAA-ATPase Vps4 to remove and recycle ESCRT-III components (Wollert et al., 2009). Accordingly, treatment of the reaction with apyrase to deplete ATP led to a reduction, but not abrogation, of PPV budding that could be restored to typical levels by addition of exogenous ATP (Fig. 7 G). Consistent with this observation, PPV release occurred rapidly upon reaction mixing, followed by further incremental release over time, suggesting that ATP acts primarily in recycling scission components (Fig. 7 G).

## Discussion

ESCRT-III components, particularly Vps20 and Snf7, can be envisaged as “peroxins”, as they are required for peroxisomal membrane biogenesis and peroxisome proliferation (Distel et al., 1996). Here, we demonstrated that ESCRT-III deletion mutants have condition-specific growth defects when grown on a non-fermentable carbon source, oleic acid (Fig. 1), the metabolism of which requires functional peroxisomes. We show that morphologically identifiable peroxisomes are absent in *vps20*Δ and *snf7*Δ cells by EM, while *did4*Δ and *vps24*Δ cells have drastically reduced numbers of peroxisomes (Fig. 2). Our experiments revealed that ESCRT-III components act as positive effectors of de novo peroxisome biogenesis, as their absence leads to fewer numbers of peroxisomes at steady-state and inhibits de novo peroxisome formation, and because they function downstream of Pex19 in this process as assessed by genetic interactions (Figs. 3 and 5). The ESCRT-III proteins Snf7 and Did4 dynamically localize to sites of de novo peroxisome biogenesis (Fig. 6), and in cell-free experiments Vps20 and Snf7 are required for the release of Pex3-containing PPVs from the ER (Fig. 7).

In the genome-wide screens that led us to ESCRT-III and its role in de novo peroxisome biogenesis, and in our experiments here (Fig. 3), import-competent peroxisomes can be found at steady-state levels in ESCRT-III deletion mutants. How can this observation be reconciled with a role for ESCRT-III in scissioning PPVs from the ER? In the case of COPI and COPII, vesicle scission can occur independently of GTP hydrolysis by simply requiring the assembly of fission-competent coated vesicles for fission to occur (Adolf et al., 2013). Thus, it is conceivable that a small proportion of budded PPVs, arrested in a fission-competent state, could be released from the ER stochastically. Following vesicle release and carrying the full complement of peroxins necessary for PPV maturation, these PPVs would become mature peroxisomes, and the growth and division cycle of peroxisome biogenesis could thereafter sustain the peroxisome population. Furthermore, as demonstrated for peroxisome fission (Motley and Hettema, 2007), we also cannot exclude a role for peroxisome inheritance factors in partially compensating for the loss of ESCRT-III. Inp2 is a membrane protein and the receptor for the motor Myo2 that actively transports peroxisomes from mother cell to bud (Fagarasanu et al., 2006; Fagarasanu et al., 2009). The presence of Inp2 in a budded vesicle at the ER could be envisaged to be sufficient to physically pull PPVs off the ER. Consistent with such a scenario is recent evidence showing preferential Inp2-directed inheritance of young peroxisomes to growing daughter cells (Kumar et al., 2017). Notwithstanding these possibilities, it is clear that peroxisomes in ESCRT-III mutants, although import-competent, are nevertheless aberrant and non-functional (Figs. 1, 2 and 3).

The formation of PPVs shares features and characteristics that are fundamental to all vesicle biogenesis events. These include cargo protein sorting and trafficking to sites of vesicle formation, the presence of adaptor molecules to guide cargo sorting and to remodel the membrane to form a budding vesicle, and scission machinery to release the vesicle to the cytosol (Spang, 2008). However, there are also fundamental differences between classical vesicle biogenesis and PPV biogenesis. Unlike the vesicles of the secretory pathway, PPVs are not coated (Titorenko and Rachubinski, 1998). PPVs also do not carry cargo in their matrix and, although it is still unknown how, must exclude components of the ER matrix. PPVs therefore share similarities with nascent lipid droplets, and similarities in PPV and lipid droplet formation, as well as shared components guiding their biogenesis, have recently come to light (Schrul and Kopito, 2016; Schuldiner and Bohnert, 2017).

Our observations are consistent with a model wherein Vps20 is recruited to sites of PPV formation, which in turn recruits and activates the polymerization of Snf7 to drive membrane scission and release of the PPV to the cytosol. Other ESCRT-III proteins like Did4 and Vps24 are also involved in PPV scission but are not essential for this process, and probably influence the dynamics of PPV formation and recruit the machinery for disassembly of ESCRT-III at the ER. Consistent with such a scenario, the *did4*Δ strain displayed severe growth defects in the presence of oleic acid and showed reduced budding of Pex3-positive PPVs from the ER (Figs. 1 and 7). Both *did4*Δ and *vps24*Δ cells showed delays in de novo peroxisome biogenesis in vivo (Fig. 3). This observation is consistent with a role for Did4 and Vps24 in recruiting the AAA-ATPase Vps4 complex to disassemble ESCRT-III polymers and with previous observations that the amount of cytosolic Snf7 is reduced in *did4*Δ cells because it is trapped in a polymerized state on membranes (Babst et al., 2002a). However, our observations may also reflect a more fundamental role for Vps24 and particularly Did4 in shaping the overall topology of the ESCRT-III polymer to be suitable for PPV scission.

ESCRT-III is typically thought to be involved in vesicle budding *away from* the cytosol (reverse-topology membrane scission) with or without other ESCRT complexes (subsets of ESCRT-0,-I and-II) (Henne et al., 2011; Schoneberg et al., 2016). Our data extend ESCRT-III function to include vesicle budding *into* the cytosol (normal-topology membrane scission). Consistent with our hypothesis, cryo-EM images of purified ESCRT-III components demonstrated that ESCRT-III can deform and stabilize membranes for normal-topology scission (McCullough et al., 2015). While normal-topology membrane scission required mammalian IST1, our observations rule out a similar role for Ist1 in peroxisome biogenesis, and instead suggest that it is the core ESCRT-III components that are capable of normal-topology membrane scission in regard to PPV biogenesis in yeast (Fig. 7). However, there may be species-specific differences in regard to ESCRT-III function in de novo peroxisome biogenesis, and it will be important to test the role of ESCRT-III in peroxisome biogenesis in other organisms. At this time, we also cannot rule out the possibility that ESCRT-III release of PPVs from the ER proceeds as a reverse-topology scission event. In this hypothetical scenario, dense tubulated matrices of ER (Nixon-Abell et al., 2016) would serve as a template for ESCRT-III to assemble as a ring through which a budded PPV could pass and be cleaved. Understanding the molecular mechanisms of ESCRT-III function in de novo peroxisome biogenesis in greater detail is undoubtedly a high priority for future research.

ESCRT-III has been implicated in unconventional protein secretion from the ER with its mode of action unclear (Curwin et al., 2016). It is interesting to note that unconventional protein secretion has previously been linked to peroxisomes (Manjithaya et al., 2010), and may more broadly share a common mechanism of egress with PPVs from the ER. Peroxisomes and ESCRT-III have also been implicated in prospore membrane formation, which requires a supply of vesicles to be delivered to and to fuse with the growing prospore membrane (Briza et al., 2002). Rather than attribute a putative metabolic role for peroxisomes in these processes, it is plausible that the non-classical vesicle secretion pathway that gives rise to peroxisomes, reliant on peroxins and ESCRT-III, is a general non-coated secretion pathway that is used more broadly by cells. The expanding role of ESCRT-III in diverse cellular activities, including the formation of peroxisomes, suggests that this evolutionarily ancient protein complex has had a much greater influence on sculpting the eukaryotic bauplan than previously realized.

## Materials and methods

### Yeast strains and plasmids

The yeast strains used in this study are listed in Table S3 and were derived from the parental strain *BY4742* or the corresponding gene deletion strain library (Invitrogen) (Giaever et al., 2002), the parental strain *W303a* (Rothstein et al., 1977), or the parental strain *R1158* (Mnaimneh et al., 2004), as described previously (Mast et al., 2016). Strains harboring genomic insertions and deletions were isolated following homologous recombination with targeted PCR fragments delivered via chemical transformation. Correct integration was verified by PCR across junctions using gene-specific primers. The following plasmids were used for PCR amplification with appropriate primers as described: pGFP/*SpHIS5* (Dilworth et al., 2001); pProtA/*SpHIS5* (Aitchison et al., 1995); pCM189/*URA3* (Gari et al., 1997); pBS34/*hph* (mCherry) (Shaner et al., 2004); and pFA6a-*nat*NT2 and pFA6a-*hph*NT1 (Janke et al., 2004). pCM189-*VPS20* was assembled by cloning a 666-bp BamHI/NotI fragment encoding full-length *VPS20* into pCM189. pCM189-*SNF7* was assembled by cloning a 723-bp BamHI/NotI fragment encoding full-length *SNF7* into pCM189.

### Yeast media and growth conditions

Yeast strains were grown in YPD medium (1% yeast extract, 2% peptone, 2% glucose) or YPBO medium (0.5% KP_i_, pH 6.0, 0.3% yeast extract, 0.5% peptone, 0.5 % Tween 40, 0.15% oleic acid), as indicated. All cultures were grown at 30°C. When marker selection was required, defined synthetic medium (SM) supplemented with 2% glucose and the necessary amino acid(s) or drug was used. Yeast media and growth conditions for specific experiments are listed below. YPB-oleate medium for use in ODELAY! was prepared as follows. A solution of 300 mg of methyl-β-cyclodextrin (Sigma)/mL and 10 μL oleic acid/mL was prepared in absolute ethanol for a total volume of 3 mL. The solution was then placed in a Rotovap for 3 h to remove the ethanol. The resulting powder was then reconstituted using ultra-pure H_2_O to a total volume of 3 mL. Meanwhile 15 mL of 1.33% agarose, 2 mL of 10 × YPB medium, and 1 mL of ultra-pure H_2_O were melted in boiling water for 18 min. Then 2 mL of the oleate-cyclodextrin solution were added to the melted medium and vortexed to mix. The YPB-oleate medium was then cast into molds and allowed to cool as described (Herricks et al., 2017a). YPB-glucose medium was prepared similarly except 2 mL of 20% glucose were added instead of the oleate-carbon source. To study peroxisome biogenesis, strains were grown overnight to saturation in YPD medium and diluted the next morning by dilution into fresh YPD medium to an OD_600_ = 0.2. Cells were then allowed to reach logarithmic phase (OD_600_ = 0.7-1) before being diluted again in YPD medium, supplemented with 4 μM DOX, and incubated for 24 h. The logarithmic phase cells were harvested by centrifugation, washed five times in YPD medium to remove DOX, inoculated into fresh YPD medium without DOX at an OD_600_ = 0.2, and cultured for an additional 24 h. At 4, 8, and 12 h, OD_600_ measurements were taken, and cultures were diluted as necessary to maintain logarithmic growth. Localization of ESCRT-III components to sites of peroxisome biogenesis used a similar growth scheme, but was performed in YPD medium containing 2 μM DOX and for 18 h.

### One-cell doubling evaluation of living arrays of yeast (ODELAY!)

Sensitive, high-density, and multiparametric analysis of cell growth was performed as described (Herricks et al., 2017a; Herricks et al., 2017b). Briefly, yeast was cultured in YPD medium in 96 well plates at 30°C overnight. Cultures were diluted to an OD_600_ = 0.09 and allowed to grow for 6 h at 30°C. The cultures were then washed in YPB medium without a carbon source, diluted to an OD_600_ = 0.02, and spotted onto YBP-oleate or YPB-glucose agarose medium. The resulting cultures were then observed using time-lapse microscopy for 48 h with 30 min intervals between images. All images were collected on a Leica DMI6000 microscope equipped with a 10× 0.3 NA lens using bright field microscopy. MATLAB scripts using the Micro-Manager interface controlled the image collection process (Edelstein et al., 2014).

Population growth rates were scored against each other using the equation:

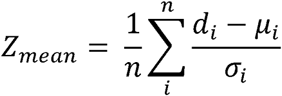

Where *d*_*i*_, is the *i*th decile of query population doubling time, *μ*_*i*_, is the mean of the ith decile of the parental strain’s doubling time, and *σ*_*i*_ is the standard deviation of the *i*th decile of the parental strain’s doubling time. The mean and standard deviation deciles (*μ*_*i*_, and *σ*_*i*_) were calculated from 16 separate *wild-type* and 8 separate deletion strain populations containing at least 200 individuals per replicate. All calculations were performed using MATLAB scripts (Herricks et al., 2017a).

### EM and quantification

Experiments were performed as described (Tam et al., 2003). Strains were cultured in YPBO medium for 8 h before being fixed and processed for EM. Image analysis to measure cell and peroxisome profiles was performed in Image J (National Institutes of Health). Stereological analysis to derive numerical densities of peroxisomes was performed as described (Weibel and Bolender, 1973).

### Fluorescence microscopy and quantification

Experiments were performed as described (Mast et al., 2016). Briefly, 10 fields of view yielding at least 100 cells per strain per time point for Gpd1-GFP labeled cells and at least 75 cells per strain per time point for Pex3-GFP/Snf7-mCherry-labeled cells or Pex3-GFP/Did4-mCherry-labeled cells were acquired in a semi-automated and randomized fashion with bright field or calcofluor white staining used to establish focus for each strain and time point. Images were acquired with a 100× 1.4 NA objective (Olympus) on a DeltaVision Elite High Resolution Microscope (GE Healthcare Life Sciences). Images were deconvolved with the manufacturer’s supplied deconvolution software (softWoRx) and an experimentally determined point spread function. Images were further processed using Imaris software (Bitplane), and object-based colocalization analysis (Bolte and Cordelieres, 2006) was performed using the “Spots” function as described previously (Mast et al., 2016). Experiments were performed in triplicate.

### Preparation of yeast whole cell lysates

Yeast whole cell lysates were prepared by denaturation in alkali solution with a reducing agent. Approximately 1.85 × 10^8^ cells (5 OD_600_ cell units) were harvested by centrifugation at 5,000 × *g* for 2 min, and resuspended in 240 μL of 1.85 M NaOH to which 7.4% (v/v) of 2-mercaptoethanol had been freshly added. The cell suspension was incubated on ice for 5 min and then mixed with an equal volume of 50% TCA by vortexing. After a 5 min incubation on ice, the precipitated protein was collected by centrifugation at 16,000 × *g* for 10 min at 4°C. The pellet was washed once with ice-cold water and resuspended in 50 μL of Magic A (1 M unbuffered Tris-HCl, 13% SDS) to which 50 μL of Magic B (30% glycerol, 200 mM DTT, 0.25% bromophenol blue) were added.

### Antibodies and antisera

Antibodies to GFP (Sigma), pA-affinity purified rabbit IgG (Mast et al., 2016), and antiserum to Kar2 (Tam et al., 2005) have been described. Antiserum to Pex19 was raised in guinea pig against full-length Pex19 lacking the carboxyl-terminal CKQQ farnesylation motif. Pex19^1-338^ was expressed as a fusion protein to glutathione-S-transferase (PEX19-GST), purified from *Escherichia coli* lysate on anti-GST Sepharose, liberated by cleavage with thrombin (removed by addition of benzamidine Sepharose), and concentrated using an Amicon Ultra centrifugal filter. Antiserum was tested for specificity for Pex19 by immunoblot of whole cell lysates from *wild-type, pex19*Δ, and *PEX19-pA* strains.

### In vitro PPV budding assay

Experiments were performed as described (Mast et al., 2016). Permeabilized yeast cells were prepared from *pex19*Δ cells expressing Pex3-pA or Pex3-GFP from an endogenous promoter. Cells were grown overnight in 1 L of YPD medium to an OD_600_ of 1. Cells were collected by centrifugation at 2,000 × *g* for 7 min at room temperature, resuspended in low glucose medium (YPD medium containing 0.1% glucose), and incubated for 30 min at 25°C with vigorous shaking. Cells were harvested by centrifugation and resuspended in spheroplast medium (1% yeast peptone, 0.1% glucose, 1.4 M sorbitol, 50 mM potassium phosphate, pH 7.5, 50 mM 2-mercaptoethanol) supplemented with yeast lytic enzyme (Zymo Research) at 1 mg/g of wet cell pellet, to a final concentration of 8 mL/g of wet cells and incubated for ^~^40 min at 37°C with gentle agitation. Spheroplasts were recovered by centrifugation and washed once in recovery medium (1% yeast peptone, 0.1 % glucose, 1 M sorbitol). Permeabilization was achieved by osmotic lysis in the presence of osmotic support. The spheroplast pellet was resuspended with ice-cold spheroplast lysis buffer (100 mM potassium acetate, 200 mM sorbitol, 20 mM HEPES-KOH, pH 7.2, 2 mM MgCl_2_) at a concentration of 5 mL/75 OD_600_ unit cell equivalents. The slurry was pipetted up and down with moderate force 5-10 times and incubated on ice for 20 min to osmotic equilibration. Permeabilized yeast cells (PYCs) were collected by centrifugation at 3,000 × *g* for 5 min at 4°C, and the supernatant was thoroughly removed. PYCs were washed twice with TBPS (115 mM potassium acetate, 2.5 mM magnesium acetate, 0.25 M sorbitol, 1× cOmplete protease inhibitor cocktail (Roche), 25 mM HEPES, pH 7.2) and resuspended in TBPS at a concentration of 25 μL/5 OD_600_ unit cell equivalents.

For yeast cytosols, spheroplasts were lysed with ice-cold 20 mM HEPES-KOH, pH 7.4, at a concentration of 210 μL per 75 OD_600_ unit cell equivalents. The slurry was pipetted 30 times with a 1 mL pipet to ensure efficient lysis. The resultant lysate was clarified by centrifugation at 3,000 × *g* for 5 min at 4°C, and the supernatant was collected and subjected to centrifugation at 100,000 × *g* for 1 h to produce a 100 k*g*S supernatant (S100) of yeast cytosol. Protein concentration was determined by a bicinchoninic acid assay (Pierce) and normalized to 4 mg/mL with addition of 10× transport buffer (250 mM HEPES-KOH, pH 7.2, 1.15 M potassium acetate, 25 mM MgCl_2_, 2.5 M sorbitol) and cOmplete protease inhibitor cocktail to 1× concentration.

Reaction conditions were as follows: 100 μL of PYCs, 100 μL of wild-type cytosol, 100 μL of a 4× ATP regenerating system (4 mM ATP, 0.4 mM GTP, 80 mM creatine phosphate, 4 U creatine phosphate kinase (Sigma)), and 100 μL of 2× TBPS were mixed on ice. The reaction was initiated by incubation at room temperature for 90 min, and chilling the samples on ice terminated the reaction. After the reaction was terminated, the PYCs were pelleted by centrifugation at 13,000 × *g* for 5 min at 4°C. To recover PPVs, the supernatant was subjected to centrifugation at 200,000 × *g* for 1 h at 4°C. The pellet was resuspended in 2× sample buffer (4% SDS, 0.15 M Tris-HCl, pH 6.8, 4 mM EDTA, 20% glycerol, 2% 2-mercaptoethanol, 0.02% bromophenol blue) before being resolved by SDS-PAGE.

To complement PPV budding defects, cytosols were mixed 1:1 at 4°C directly before addition to the reaction. For experiments with apyrase, PYCs and cytosols were incubated separately with 1 U of apyrase (Sigma) and resuspended in reaction buffer (25 mM HEPES-KOH, pH 7.2, 115 mM potassium acetate, 2.5 mM MgCl_2_, 250 mM sorbitol) for 20 min before starting the reaction. For experiments in which exogenous ATP was added back, only apyrase-treated PYCs that had been washed once in reaction buffer were used. Alphaview (ProteinSimple) was used to quantify the chemiluminescence signal with values normalized between the negative *pex19*Δ PYCs only control (set to 0), and the positive *wild-type* cytosol plus ATP control (set to 100). Experiments were performed in triplicate.

## Online supplemental material

Fig. S1 contains supporting data to show additional electron micrographs of the ESCRT-III deletion mutants revealing peroxisome morphology defects. Fig. S2 contains supporting data to show the budding efficiencies of PPVs from reactions containing cytosols isolated from ESCRT deletion strains. Fig. S3 contains supporting data to show the levels of Pex19 in ESCRT deletion mutants. Table S1 contains supplemental information summarizing the results of ESCRT deletion mutants from previous genomic screens for peroxisome function. Table S2 contains supplemental information on the normalized *Z*-scores for all strains measured by ODELAY!. Table S3 contains supplemental information on the genetic background of the yeast strains used in this study, as well as their origin of derivation.

## Acknowledgments

We thank Alexis Kaushansky, Sanjeev Kumar, Maxwell Neal, and the Aitchison laboratory for helpful discussion and feedback on the manuscript. We gratefully acknowledge the technical assistance of Elena Savidov.

This work was supported by a Foundation Grant from the Canadian Institutes of Health Research (CIHR) to R.A.R. and by grants P50 GM076547 and P41 GM109824 from the National Institutes of Health to J.D.A. F.D.M is a postdoctoral fellow of the CIHR. The support of Ian Tolmie and Jennifer Black is gratefully acknowledged, and this paper is dedicated to the memory their son, Dane Tolmie.

The authors declare no competing financial interests.

## Author contributions

F.D.M., R.A.R, and J.D.A. designed the experiments, analyzed the results and wrote the manuscript. F.D.M. performed the experiments. R.A.R performed the EM experiments. T.H. performed the ODELAY! experiments and analysis. K.M.S and L.R.M assisted with the experiments. R.A.S contributed to the experimental design and analysis. All authors read and commented on the manuscript.

## Abbreviations

DOX: doxycycline
DRP: dynamin-related protein
ESCRT: endosomal sorting complexes required for transport
PBD: peroxisome biogenesis disorder
*PEX*: gene encoding a peroxin
PMP: peroxisomal membrane protein
PPV: preperoxisomal vesicle

## References

Adolf, F., A. Herrmann, A. Hellwig, R. Beck, B. Brugger, and F.T. Wieland. 2013. Scission of COPI and COPII vesicles is independent of GTP hydrolysis. Traffic. 14:922–932.

Agrawal, G., S.N. Fassas, Z.J. Xia, and S. Subramani. 2016. Distinct requirements for intra-ER sorting and budding of peroxisomal membrane proteins from the ER. J. Cell Biol. 212:335–348.

Agrawal, G., S. Joshi, and S. Subramani. 2011. Cell-free sorting of peroxisomal membrane proteins from the endoplasmic reticulum. Proc. Natl. Acad. Sci. USA. 108:9113–9118.

Agrawal, G., H.H. Shang, Z.J. Xia, and S. Subramani. 2017. Functional regions of the peroxin Pex19 necessary for peroxisome biogenesis. J. Biol. Chem. 292:11547–11560.

Agrawal, G., and S. Subramani. 2016. De novo peroxisome biogenesis: Evolving concepts and conundrums. Biochim. Biophys. Acta. 1863:892–901.

Aitchison, J.D., G. Blobel, and M.P. Rout. 1995. Nup120p: a yeast nucleoporin required for NPC distribution and mRNA transport. J. Cell Biol. 131:1659–1675.

Babst, M., D.J. Katzmann, E.J. Estepa-Sabal, T. Meerloo, and S.D. Emr. 2002a. ESCRT-III: an endosome-associated heterooligomeric protein complex required for MVB sorting. Dev. Cell. 3:271–282.

Babst, M., D.J. Katzmann, W.B. Snyder, B. Wendland, and S.D. Emr. 2002b. Endosome-associated complex, ESCRT-II, recruits transport machinery for protein sorting at the multivesicular body. Dev. Cell. 3:283–289.

Babst, M., T.K. Sato, L.M. Banta, and S.D. Emr. 1997. Endosomal transport function in yeast requires a novel AAA-type ATPase, Vps4p. EMBO J. 16:1820–1831.

Bauer, S.T., K.E. McQueeney, T. Patel, and M.T. Morris. 2017. Localization of a trypanosome peroxin to the endoplasmic reticulum. J. Eukaryot. Microbiol. 64:97–105.

Bolte, S., and F.P. Cordelieres. 2006. A guided tour into subcellular colocalization analysis in light microscopy. J. Microsc. 224:213–232.

Briza, P., E. Bogengruber, A. Thur, M. Rutzler, M. Munsterkotter, I.W. Dawes, and M. Breitenbach. 2002. Systematic analysis of sporulation phenotypes in 624 non-lethal homozygous deletion strains of Saccharomyces cerevisiae. Yeast. 19:403–422.

Carlton, J.G., and J. Martin-Serrano. 2007. Parallels between cytokinesis and retroviral budding: a role for the ESCRT machinery. Science. 316:1908–1912.

Colasante, C., J. Chen, B. Ahlemeyer, and E. Baumgart-Vogt. 2015. Peroxisomes in cardiomyocytes and the peroxisome / peroxisome proliferator-activated receptor-loop. Thromb. Haemost. 113:452–463.

Costanzo, M., B. VanderSluis, E.N. Koch, A. Baryshnikova, C. Pons, G. Tan, W. Wang, M. Usaj, J. Hanchard, S.D. Lee, et al. 2016. A global genetic interaction network maps a wiring diagram of cellular function. Science. 353.

Curwin, A.J., N. Brouwers, Y.A.M. Alonso, D. Teis, G. Turacchio, S. Parashuraman, P. Ronchi, and V. Malhotra. 2016. ESCRT-III drives the final stages of CUPS maturation for unconventional protein secretion. eLife. 5.

David, C., J. Koch, S. Oeljeklaus, A. Laernsack, S. Melchior, S. Wiese, A. Schummer, R. Erdmann, B. Warscheid, and C. Brocard. 2013. A combined approach of quantitative interaction proteomics and live-cell imaging reveals a regulatory role for endoplasmic reticulum (ER) reticulon homology proteins in peroxisome biogenesis. Mol. Cell. Proteomics. 12:2408–2425.

De Craemer, D., M. Pauwels, and F. Roels. 1993. Peroxisomes in cirrhosis of the human liver: a cytochemical, ultrastructural and quantitative study. Hepatology. 17:404–410.

Delmaghani, S., J. Defourny, A. Aghaie, M. Beurg, D. Dulon, N. Thelen, I. Perfettini, T. Zelles, M. Aller, A. Meyer, et al. 2015. Hypervulnerability to sound exposure through impaired adaptive proliferation of peroxisomes. Cell. 163:894–906.

Di Cara, F., A. Sheshachalam, N.E. Braverman, R.A. Rachubinski, and A.J. Simmonds. 2017. Peroxisome-mediated metabolism is required for immune response to microbial infection. Immunity. 47:93–106.

Dilworth, D.J., A. Suprapto, J.C. Padovan, B.T. Chait, R.W. Wozniak, M.P. Rout, and J.D. Aitchison. 2001. Nup2p dynamically associates with the distal regions of the yeast nuclear pore complex. J. Cell Biol. 153:1465–1478.

Distel, B., R. Erdmann, S.J. Gould, G. Blobel, D.I. Crane, J.M. Cregg, G. Dodt, Y. Fujiki, J.M. Goodman, W.W. Just, et al. 1996. A unified nomenclature for peroxisome biogenesis factors. J. Cell Biol. 135:1–3.

Dixit, E., S. Boulant, Y. Zhang, A.S. Lee, C. Odendall, B. Shum, N. Hacohen, Z.J. Chen, S.P. Whelan, M. Fransen, et al. 2010. Peroxisomes are signaling platforms for antiviral innate immunity. Cell. 141:668–681.

Dixon, S.J., M. Costanzo, A. Baryshnikova, B. Andrews, and C. Boone. 2009. Systematic mapping of genetic interaction networks. Ann. Rev. Genet. 43:601–625.

Drees, B.L., V. Thorsson, G.W. Carter, A.W. Rives, M.Z. Raymond, I. Avila-Campillo, P. Shannon, and T. Galitski. 2005. Derivation of genetic interaction networks from quantitative phenotype data. Genome Biol. 6:R38.

Duden, R., L. Kajikawa, L. Wuestehube, and R. Schekman. 1998. ɛ-COP is a structural component of coatomer that functions to stabilize α-COP. EMBO J. 17:985–995.

Edelstein, A.D., M.A. Tsuchida, N. Amodaj, H. Pinkard, R.D. Vale, and N. Stuurman. 2014. Advanced methods of microscope control using μManager software. J. Biol. Methods. 1:e10.

Fagarasanu, A., M. Fagarasanu, G.A. Eitzen, J.D. Aitchison, and R.A. Rachubinski. 2006. The peroxisomal membrane protein Inp2p is the peroxisome-specific receptor for the myosin V motor Myo2p of Saccharomyces cerevisiae. Dev. Cell. 10:587–600.

Fagarasanu, A., F.D. Mast, B. Knoblach, Y. Jin, M.J. Brunner, M.R. Logan, J.N. Glover, G.A. Eitzen, J.D. Aitchison, L.S. Weisman, et al. 2009. Myosin-driven peroxisome partitioning in S. cerevisiae. J. Cell Biol. 186:541–554.

Fakieh, M.H., P.J. Drake, J. Lacey, J.M. Munck, A.M. Motley, and E.H. Hettema. 2013. Intra-ER sorting of the peroxisomal membrane protein Pex3 relies on its luminal domain. Biol. Open. 2:829–837.

Farré, J.C., K. Carolino, O.V. Stasyk, O.G. Stasyk, Z. Hodzic, G. Agrawal, A. Till, M. Proietto, J. Cregg, A.A. Sibirny, et al. 2017. A new yeast peroxin, Pex36, a functional homolog of mammalian PEX16, functions in the ER-to-peroxisome traffic of peroxisomal membrane proteins. J. Mol. Biol. 429:3743–3762.

Gari, E., L. Piedrafita, M. Aldea, and E. Herrero. 1997. A set of vectors with a tetracycline-regulatable promoter system for modulated gene expression in Saccharomyces cerevisiae. Yeast. 13:837–848.

Garrus, J.E., U.K. von Schwedler, O.W. Pornillos, S.G. Morham, K.H. Zavitz, H.E. Wang, D.A. Wettstein, K.M. Stray, M. Cote, R.L. Rich, et al. 2001. Tsg101 and the vacuolar protein sorting pathway are essential for HIV-1 budding. Cell. 107:55–65.

Giaever, G., A.M. Chu, L. Ni, C. Connelly, L. Riles, S. Véronneau, S. Dow, A. Lucau-Danila, K. Anderson, B. André, et al. 2002. Functional profiling of the Saccharomyces cerevisiae genome. Nature. 418:387–391.

Guizetti, J., L. Schermelleh, J. Mantler, S. Maar, I. Poser, H. Leonhardt, T. Muller-Reichert, and D.W. Gerlich. 2011. Cortical constriction during abscission involves helices of ESCRT-III-dependent filaments. Science. 331:1616–1620.

Guther, M.L., M.D. Urbaniak, A. Tavendale, A. Prescott, and M.A. Ferguson. 2014. High-confidence glycosome proteome for procyclic form Trypanosoma brucei by epitope-tag organelle enrichment and SILAC proteomics. J. Proteome Res. 13:2796–2806.

Halbach, A., C. Landgraf, S. Lorenzen, K. Rosenkranz, R. Volkmer-Engert, R. Erdmann, and H. Rottensteiner. 2006. Targeting of the tail-anchored peroxisomal membrane proteins PEX26 and PEX15 occurs through C-terminal PEX19-binding sites. J. Cell Sci. 119:2508–2517.

Henne, W.M., N.J. Buchkovich, and S.D. Emr. 2011. The ESCRT pathway. Dev. Cell. 21:77–91.

Herricks, T., D.J. Dilworth, F.D. Mast, S. Li, J.J. Smith, A.V. Ratushny, and J.D. Aitchison. 2017a. One-cell doubling evaluation by living arrays of yeast, ODELAY! G3 (Bethesda). 7:279–288.

Herricks, T., F.D. Mast, S. Li, and J.D. Aitchison. 2017b. ODELAY: A large-scale method for multi-parameter quantification of yeast growth. J. Vis. Exp., e55879.

Hettema, E.H., W. Girzalsky, M. van Den Berg, R. Erdmann, and B. Distel. 2000. Saccharomyces cerevisiae Pex3p and Pex19p are required for proper localization and stability of peroxisomal membrane proteins. EMBO J. 19:223–233.

Hoepfner, D., D. Schildknegt, I. Braakman, P. Philippsen, and H.F. Tabak. 2005. Contribution of the endoplasmic reticulum to peroxisome formation. Cell. 122:85–95.

Hu, J., A. Baker, B. Bartel, N. Linka, R.T. Mullen, S. Reumann, and B.K. Zolman. 2012. Plant peroxisomes: biogenesis and function. Plant Cell. 24:2279–2303.

Igual, J.C., E. Matallana, C. Gonzalez-Bosch, L. Franco, and J.E. Perez-Ortin. 1991. A new glucose-repressible gene identified from the analysis of chromatin structure in deletion mutants of yeast SUC2 locus. Yeast. 7:379–389.

Jan, C.H., C.C. Williams, and J.S. Weissman. 2014. Principles of ER cotranslational translocation revealed by proximity-specific ribosome profiling. Science. 346:1257521.

Janke, C., M.M. Magiera, N. Rathfelder, C. Taxis, S. Reber, H. Maekawa, A. Moreno-Borchart, G. Doenges, E. Schwob, E. Schiebel, et al. 2004. A versatile toolbox for PCR-based tagging of yeast genes: new fluorescent proteins, more markers and promoter substitution cassettes. Yeast. 21:947–962.

Jimenez, A.J., P. Maiuri, J. Lafaurie-Janvore, S. Divoux, M. Piel, and F. Perez. 2014. ESCRT machinery is required for plasma membrane repair. Science. 343:1247136.

Joshi, A.S., X. Huang, V. Choudhary, T.P. Levine, J. Hu, and W.A. Prinz. 2016. A family of membrane-shaping proteins at ER subdomains regulates pre-peroxisomal vesicle biogenesis. J. Cell Biol. 215:515–529.

Jung, S., M. Marelli, R.A. Rachubinski, D.R. Goodlett, and J.D. Aitchison. 2010. Dynamic changes in the subcellular distribution of Gpd1p in response to cell stress. J. Biol. Chem. 285:6739–6749.

Kaewsapsak, P., D.M. Shechner, W. Mallard, J.L. Rinn, and A.Y. Ting. 2017. Live-cell mapping of organelle-associated RNAs via proximity biotinylation combined with protein-RNA crosslinking. eLife. 6:e29224.

Katzmann, D.J., M. Babst, and S.D. Emr. 2001. Ubiquitin-dependent sorting into the multivesicular body pathway requires the function of a conserved endosomal protein sorting complex, ESCRT-I. Cell. 106:145–155.

Katzmann, D.J., C.J. Stefan, M. Babst, and S.D. Emr. 2003. Vps27 recruits ESCRT machinery to endosomes during MVB sorting. J. Cell Biol. 162:413–423.

Kao, Y.T., K.L. Gonzalez, and B. Bartel. 2018. Peroxisome function, biogenesis, and dynamics in plants. Plant Physiol. 176:162–177.

Kim, P.K., R.T. Mullen, U. Schumann, and J. Lippincott-Schwartz. 2006. The origin and maintenance of mammalian peroxisomes involves a de novo PEX16-dependent pathway from the ER. J. Cell Biol. 173:521–532.

Kumar, S., R. de Boer, and I. van der Klei. 2018. Yeast cells contain a heterogeneous population of peroxisomes that segregate asymmetrically during cell division. J. Cell Sci. In press.

Lam, S.K., N. Yoda, and R. Schekman. 2010. A vesicle carrier that mediates peroxisome protein traffic from the endoplasmic reticulum. Proc. Natl. Acad. Sci. USA. 107:21523–21528.

Lee, J.A., A. Beigneux, S.T. Ahmad, S.G. Young, and F.B. Gao. 2007. ESCRT-III dysfunction causes autophagosome accumulation and neurodegeneration. Curr. Biol. 17:1561–1567.

Li, X., E. Baumgart, J.C. Morrell, G. Jimenez-Sanchez, D. Valle, and S.J. Gould. 2002. PEX11β deficiency is lethal and impairs neuronal migration but does not abrogate peroxisome function. Mol. Cell. Biol. 22:4358–4365.

Loncle, N., M. Agromayor, J. Martin-Serrano, and D.W. Williams. 2015. An ESCRT module is required for neuron pruning. Sci. Rep. 5:8461.

Mani, R., R.P. St. Onge, J.L. Hartman IV, G. Giaever, and F.P. Roth. 2008. Defining genetic interaction. Proc. Natl. Acad. Sci. USA. 105:3461–3466.

Manjithaya, R., C. Anjard, W.F. Loomis, and S. Subramani. 2010. Unconventional secretion of Pichia pastoris Acb1 is dependent on GRASP protein, peroxisomal functions, and autophagosome formation. J. Cell Biol. 188:537–546.

Mast, F.D., A. Fagarasanu, B. Knoblach, and R.A. Rachubinski. 2010. Peroxisome biogenesis: something old, something new, something borrowed. Physiology. 25:347–356.

Mast, F.D., A. Jamakhandi, R.A. Saleem, D.J. Dilworth, R.S. Rogers, R.A. Rachubinski, and J.D. Aitchison. 2016. Peroxins Pex30 and Pex29 dynamically associate with reticulons to regulate peroxisome biogenesis from the endoplasmic reticulum. J. Biol. Chem. 291:154–8015427.

Mast, F.D., R.A. Rachubinski, and J.D. Aitchison. 2015. Signaling dynamics and peroxisomes. Curr. Opin. Cell Biol. 35:131–136.

Mayerhofer, P.U., M. Bano-Polo, I. Mingarro, and A.E. Johnson. 2016. Human peroxin PEX3 is co-translationally Integrated into the ER and exits the ER in budding vesicles. Traffic. 17:117–130.

McCullough, J., A.K. Clippinger, N. Talledge, M.L. Skowyra, M.G. Saunders, T.V. Naismith, L.A. Colf, P. Afonine, C. Arthur, W.I. Sundquist, et al. 2015. Structure and membrane remodeling activity of ESCRT-III helical polymers. Science. 350:1548–1551.

Meinecke, M., C. Cizmowski, W. Schliebs, V. Kruger, S. Beck, R. Wagner, and R. Erdmann. 2010. The peroxisomal importomer constitutes a large and highly dynamic pore. Nat. Cell Biol. 12:273–277.

Mnaimneh, S., A.P. Davierwala, J. Haynes, J. Moffat, W.T. Peng, W. Zhang, X. Yang, J. Pootoolal, G. Chua, A. Lopez, et al. 2004. Exploration of essential gene functions via titratable promoter alleles. Cell. 118:31–44.

Motley, A.M., P.C. Galvin, L. Ekal, J.M. Nuttall, and E.H. Hettema. 2015. Reevaluation of the role of Pex1 and dynamin-related proteins in peroxisome membrane biogenesis. J. Cell Biol. 211:1041–1056.

Motley, A.M., and E.H. Hettema. 2007. Yeast peroxisomes multiply by growth and division. J. Cell Biol. 178:399–410.

Motley, A.M., G.P. Ward, and E.H. Hettema. 2008. Dnm1p-dependent peroxisome fission requires Caf4p, Mdv1p and Fis1p. J. Cell Sci. 121:1633–1640.

Nichols, B.J., C. Ungermann, H.R. Pelham, W.T. Wickner, and A. Haas. 1997. Homotypic vacuolar fusion mediated by t-and v-SNAREs. Nature. 387:199–202.

Nixon-Abell, J., C.J. Obara, A.V. Weigel, D. Li, W.R. Legant, C.S. Xu, H.A. Pasolli, K. Harvey, H.F. Hess, E. Betzig, et al. 2016. Increased spatiotemporal resolution reveals highly dynamic dense tubular matrices in the peripheral ER. Science. 354:aaf3928.

Odendall, C., E. Dixit, F. Stavru, H. Bierne, K.M. Franz, A.F. Durbin, S. Boulant, L. Gehrke, P. Cossart, and J.C. Kagan. 2014. Diverse intracellular pathogens activate type III interferon expression from peroxisomes. Nat. Immunol. 15:717–726.

Olmos, Y., L. Hodgson, J. Mantell, P. Verkade, and J.G. Carlton. 2015. ESCRT-III controls nuclear envelope reformation. Nature. 522:236–239.

Perry, R.J., F.D. Mast, and R.A. Rachubinski. 2009. Endoplasmic reticulum-associated secretory proteins Sec20p, Sec39p, and Dsl1p are involved in peroxisome biogenesis. Euk. Cell. 8:830–843.

Reumann, S., and B. Bartel. 2016. Plant peroxisomes: recent discoveries in functional complexity, organelle homeostasis, and morphological dynamics. Curr. Opin. Plant Biol. 34:17–26.

Reumann, S., and A.P. Weber. 2006. Plant peroxisomes respire in the light: some gaps of the photorespiratory C2 cycle have become filled 0 others remain. Biochem. Biophys. Acta. 1763:1496–1510.

Rothstein, R.J., R.E. Esposito, and M.S. Esposito. 1977. The effect of ochre suppression on meiosis and ascospore formation in Saccharomyces. Genetics. 85:35–54.

Rusten, T.E., T. Vaccari, K. Lindmo, L.M. Rodahl, I.P. Nezis, C. Sem-Jacobsen, F. Wendler, J.P. Vincent, A. Brech, D. Bilder, et al. 2007. ESCRTs and Fab1 regulate distinct steps of autophagy. Curr. Biol. 17:1817–1825.

Saleem, R.A., B. Knoblach, F.D. Mast, J.J. Smith, J. Boyle, C.M. Dobson, R. Long-O’Donnell, R.A. Rachubinski, and J.D. Aitchison. 2008. Genome-wide analysis of signaling networks regulating fatty acid-induced gene expression and organelle biogenesis. J. Cell Biol. 181:281–292.

Saleem, R.A., R. Long-O’Donnell, D.J. Dilworth, A.M. Armstrong, A.P. Jamakhandi, Y. Wan, T.A. Knijnenburg, A. Niemisto, J. Boyle, R.A. Rachubinski, et al. 2010. Genome-wide analysis of effectors of peroxisome biogenesis. PLOS ONE. 5:e11953.

Sato, K., and A. Nakano. 2002. Emp47p and its close homolog Emp46p have a tyrosine-containing endoplasmic reticulum exit signal and function in glycoprotein secretion in Saccharomyces cerevisiae. Mol. Biol. Cell. 13:2518–2532.

Scheffer, L.L., S.C. Sreetama, N. Sharma, S. Medikayala, K.J. Brown, A. Defour, and J.K. Jaiswal. 2014. Mechanism of Ca^2+^-triggered ESCRT assembly and regulation of cell membrane repair. Nat. Commun. 5:5646.

Schoneberg, J., I.H. Lee, J.H. Iwasa, and J.H. Hurley. 2016. Reverse-topology membrane scission by the ESCRT proteins. Nat. Rev. Mol. Cell Biol. 18:5–17.

Schrader, M., J.L. Costello, L.F. Godinho, A.S. Azadi, and M. Islinger. 2016. Proliferation and fission of peroxisomes - An update. Biochim. Biophys. Acta. 1863:971–983.

Schrul, B., and R.R. Kopito. 2016. Peroxin-dependent targeting of a lipid-droplet-destined membrane protein to ER subdomains. Nat. Cell Biol. 18:740–751.

Schuldiner, M., and M. Bohnert. 2017. A different kind of love - lipid droplet contact sites. Biochim. Biophys. Acta. 1862:1188–1196.

Schuldiner, M., S.R. Collins, N.J. Thompson, V. Denic, A. Bhamidipati, T. Punna, J. Ihmels, B. Andrews, C. Boone, J.F. Greenblatt, et al. 2005. Exploration of the function and organization of the yeast early secretory pathway through an epistatic miniarray profile. Cell. 123:507–519.

Schuldiner, M., J. Metz, V. Schmid, V. Denic, M. Rakwalska, H.D. Schmitt, B. Schwappach, and J.S. Weissman. 2008. The GET complex mediates insertion of tail-anchored proteins into the ER membrane. Cell. 134:634–645.

Shaner, N.C., R.E. Campbell, P.A. Steinbach, B.N. Giepmans, A.E. Palmer, and R.Y. Tsien. 2004. Improved monomeric red, orange and yellow fluorescent proteins derived from Discosoma sp. red fluorescent protein. Nat. Biotechnol. 22:1567–1572.

Smith, J.J., and J.D. Aitchison. 2013. Peroxisomes take shape. Nat. Rev. Mol. Cell Biol. 14:803–817.

Smith, J.J., M. Marelli, R.H. Christmas, F.J. Vizeacoumar, D.J. Dilworth, T. Ideker, T. Galitski, K. Dimitrov, R.A. Rachubinski, and J.D. Aitchison. 2002. Transcriptome profiling to identify genes involved in peroxisome assembly and function. J. Cell Biol. 158:259–271.

Smith, J.J., Y. Sydorskyy, M. Marelli, D. Hwang, H. Bolouri, R.A. Rachubinski, and J.D. Aitchison. 2006. Expression and functional profiling reveal distinct gene classes involved in fatty acid metabolism. Mol. Syst. Biol. 2:2006 0009.

South, S.T., K.A. Sacksteder, X. Li, Y. Liu, and S.J. Gould. 2000. Inhibitors of COPI and COPII do not block PEX3-mediated peroxisome synthesis. J. Cell Biol. 149:1345–1360.

Spang, A. 2008. The life cycle of a transport vesicle. Cell Mol. Life Sci. 65:2781–2789.

Tam, Y.Y.C., A. Fagarasanu, M. Fagarasanu, and R.A. Rachubinski. 2005. Pex3p initiates the formation of a preperoxisomal compartment from a subdomain of the endoplasmic reticulum in Saccharomyces cerevisiae. J. Biol. Chem. 280:34933–34939.

Tam, Y.Y.C, J.C. Torres-Guzman, F.J. Vizeacoumar, J.J. Smith, M. Marelli, J.D. Aitchison, and R.A. Rachubinski. 2003. Pex11-related proteins in peroxisome dynamics: a role for the novel peroxin Pex27p in controlling peroxisome size and number in Saccharomyces cerevisiae. Mol. Biol. Cell. 14:4089–4102.

Tang, S., W.M. Henne, P.P. Borbat, N.J. Buchkovich, J.H. Freed, Y. Mao, J.C. Fromme, and S.D. Emr. 2015. Structural basis for activation, assembly and membrane binding of ESCRT-III Snf7 filaments. eLife. 4:e12548.

Thoms, S., I. Harms, K.U. Kalies, and J. Gartner. 2012. Peroxisome formation requires the endoplasmic reticulum channel protein Sec61. Traffic. 13:599–609.

Titorenko, V.I., H. Chan, and R.A. Rachubinski. 2000. Fusion of small peroxisomal vesicles in vitro reconstructs an early step in the in vivo multistep peroxisome assembly pathway of Yarrowia lipolytica. J. Cell Biol. 148:29–44.

Titorenko, V.I., and R.A. Rachubinski. 1998. Mutants of the yeast Yarrowia lipolytica defective in protein exit from the endoplasmic reticulum are also defective in peroxisome biogenesis. Mol. Cell. Biol. 18:2789–2803.

Titorenko, V.I., and R.A. Rachubinski. 2000. Peroxisomal membrane fusion requires two AAA family ATPases, Pex1p and Pex6p. J. Cell Biol. 150:881–886.

Tripathi, D.N., and C.L. Walker. 2016. The peroxisome as a cell signaling organelle. Curr. Opin. Cell Biol. 39:109–112.

van der Zand, A., I. Braakman, and H.F. Tabak. 2010. Peroxisomal membrane proteins insert into the endoplasmic reticulum. Mol. Biol. Cell. 21:2057–2065.

van der Zand, A., J. Gent, I. Braakman, and H.F. Tabak. 2012. Biochemically distinct vesicles from the endoplasmic reticulum fuse to form peroxisomes. Cell. 149:397–409.

Vietri, M., K.O. Schink, C. Campsteijn, C.S. Wegner, S.W. Schultz, L. Christ, S.B. Thoresen, A. Brech, C. Raiborg, and H. Stenmark. 2015. Spastin and ESCRT-III coordinate mitotic spindle disassembly and nuclear envelope sealing. Nature. 522:231–235.

Webster, B.M., P. Colombi, J. Jager, and C.P. Lusk. 2014. Surveillance of nuclear pore complex assembly by ESCRT-III/Vps4. Cell. 159:388–401.

Weibel, E.R., and R.P. Bolender. 1973. Stereological techniques for electron microscopic morphometry. In Principles and Techniques of Electron Microscopy. M.A. Hayat, editor. Van Nostrand Reinhold, New York, NY. 237–296.

Weller, S., S.J. Gould, and D. Valle. 2003. Peroxisome biogenesis disorders. Ann. Rev. Genomics Hum. Genet. 4:165–211.

Wollert, T., C. Wunder, J. Lippincott-Schwartz, and J.H. Hurley. 2009. Membrane scission by the ESCRT-III complex. Nature. 458:172–177.

Young, J.M., J.W. Nelson, J. Cheng, W. Zhang, S. Mader, C.M. Davis, R.S. Morrison, and N.J. Alkayed. 2015. Peroxisomal biogenesis in ischemic brain. Antioxid. Redox Signal. 22:109–120.

Zhang, J., J. Kim, A. Alexander, S. Cai, D.N. Tripathi, R. Dere, A.R. Tee, J. Tait-Mulder, A. Di Nardo, J.M. Han, et al. 2013. A tuberous sclerosis complex signalling node at the peroxisome regulates mTORC1 and autophagy in response to ROS. Nat. Cell Biol. 15:1186–1196.

Zhang, J., D.N. Tripathi, J. Jing, A. Alexander, J. Kim, R.T. Powell, R. Dere, J. Tait-Mulder, J.H. Lee, T.T. Paull, et al. 2015. ATM functions at the peroxisome to induce pexophagy in response to ROS. Nat. Cell Biol. 17:1259–1269.

